# Structure and assembly of a *Clostridioides difficile* spore polar appendage

**DOI:** 10.1101/468637

**Authors:** Wilson Antunes, Fátima C. Pereira, Carolina Feliciano, Laure Saujet, Tiago dos Vultos, Evelyne Couture-Tosi, Severine Péchiné, Jean-François Bruxelle, Claire Janoir, Luís V. Melo, Patrícia Brito, Isabelle Martin-Verstraete, Mónica Serrano, Bruno Dupuy, Adriano O. Henriques

## Abstract

*Clostridioides difficile*, a strict anaerobic spore-former, is the main cause of nosocomial disease associated to antibiotic therapy in adults and a growing concern in the community. Spores are the main infectious, persistence and transmission vehicle. Spore germination occurs in the intestine and the resulting vegetative cells will produce the toxins responsible for the disease symptoms, and spores. During sporulation, a wild type population bifurcates into two main spore morphotypes, with or without a thick exosporium. We show that this bifurcation extends to the formation of spores with a robust polar appendage or spores with a short appendage or that lack this structure. The cysteine-rich CdeM protein localizes to the appendage and around the entire surface of the spore, and is a major structural component of the exosporium, which we show is continuous with the appendage. In a *CdeM* mutant, when present, the polar appendage is short and disorganized. We show that wild type and *cdeM* spores with a short or no appendage germinate poorly in response to taurocholate, compared to those with an appendage. *cdeM* spores of the two types, however, germinate faster than their wild type counterparts. Thus, while the absence of CdeM may increase the permeability of spores to taurocholate, proper assembly of the appendage is also important for germination. Consistent with an overall enhancement of germination, a *cdeM* mutant shows increased virulence in a hamster model of disease. For a wild type population, spores with a short or no appendage germinate slower than the appendage-bearing spores. Differences in transmission, persistence and disease severity may result, in part, from their proportion in a spore population.

## Introduction

Infection by *Clostridium difficile* (CDI), recently renamed *Clostridioides difficile* (Lawson *et al.*, 2016, Yutin & Galperin, 2013), is the cause of an intestinal disease mediated by two toxins, TcdA and TcdB (Aktories *et al.*, 2017, Chandrasekaran & Lacy, 2017). Symptoms of CDI can range from asymptomatic colonization or mild diarrhea, to more severe conditions that can lead to death (Rupnik *et al.*, 2009, Smits *et al.*, 2016). The last two decades have witnessed the emergence and spreading of epidemic strains which have caused outbreaks associated with more severe disease symptoms, morbidity, mortality and disease recurrence rates (Smits *et al.*, 2016). This has led to the recognition of *C. difficile* as a main nosocomial enteric pathogen and its recognition as an urgent threat (Kociolek & Gerding, 2016, Rupnik *et al.*, 2009, Smits *et al.*, 2016). Health care-associated CDI develops in hospitalized patients undergoing antibiotic treatment because *C. difficile* can colonize the gut if the protective effect of the intestinal microbiota is lifted (Rupnik *et al.*, 2009, Smits *et al.*, 2016). Recent changes in the epidemiology of this pathogen, together with its emergence at the community level and the risk of widespread zoonotic transmission, adds to the problem (Dubberke & Olsen, 2012, Isidro *et al.*, 2018, Jones *et al.*, 2013, Lessa *et al.*, 2015, Rupnik *et al.*, 2009, Smits *et al.*, 2016).

*C. difficile* is an obligate anaerobe, and has the ability to form endospores (spores for simplicity) which are extremely resilient and hard to eradicate (Setlow, 2014). The oxygen-resistant spores are the infectious and transmissible form of the organism; their persistence within the host or in environmental reservoirs is also linked to disease recurrence (Deakin *et al.*, 2012, Paredes-Sabja & Sarker, 2012). Sporulation begins with a polar division that creates a larger mother cell and a smaller forespore, or future spore (Fig. 1A). Following polar division, cell type-specific gene expression is controlled by four RNA polymerase sigma subunits; σ^F^ and σ^E^ drive the early stages of development in the mother cell and in the forespore, and are replaced at later stages by σ^G^ and σ^K^ (Driks & Eichenberger, 2016, Fimlaid *et al.*, 2013, Fimlaid & Shen, 2015, Henriques & Moran, 2007, McKenney *et al.*, 2013, Pereira *et al.*, 2013, Saujet *et al.*, 2013) (Fig. 1A). The genes coding for these sigma factors are part of a genomic signature of sporulation (Abecasis *et al.*, 2013). Soon after polar division, the mother cell engulfs the forespore, and at late stages in development, it directs assembly of a thick layer of a modified form of peptidoglycan, called the cortex, essential to maintain spore dormancy, and of several multiprotein layers which encase the cortex (Fig. 1A). The components of these proteinaceous layers are produced under the direct control of σ^E^ and σ^K^ (Driks & Eichenberger, 2016, Henriques & Moran, 2007, McKenney *et al.*, 2013) (Fig. 1A).

**Figure 1.**
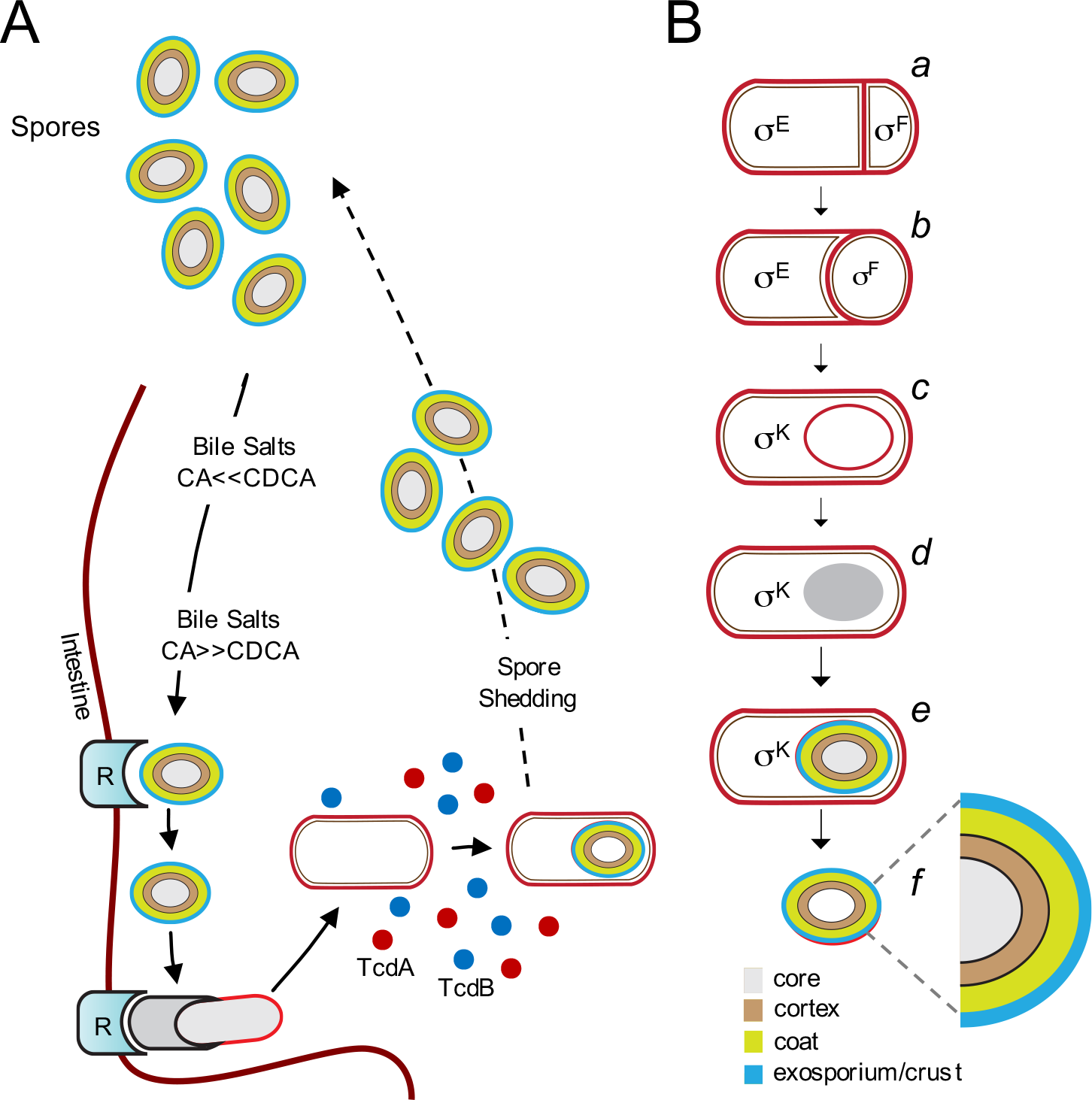
Sporulation and infection by *C. difficile.* **A**: Schematic representation of the *C. difficile* infectious cycle with emphasis on the role of the spore. Infection starts with spore ingestion. Spores germinate in the intestine, in response to bile salts in the form of cholate, which are more abundant than inhibitory deoxycholate salts (CA<<CDCA). Spore germination seems to occur in close contact with epithelial cells. “R” is a putative spore receptor at the surface of epithelial cells. Following spore germination and outgrowth, a population of vegetative cells is established that produces the TcdA and TcdB toxins and spores. Shedding of spores from the infected organism completes the cycle. **B**: Stages in spore morphogenesis: *a*, asymmetric septation; *b*, an intermediate stage in engulfment; *c*, engulfment completion; *d*, phase grey spores; *e*, phase bright spores; *f*, free spores. The main spore layers are represented as follows: yellow, core; brown, cortex; blue, coats; red, crust or exosporium. The four cell-type specific sigma factors that control spore morphogenesis and their windows of activity with respect to the stages in morphogenesis are also indicated.

The structure and composition of the spore surface varies among species. In *B. subtilis*, the outermost spore structures are a coat and a crust (Driks & Eichenberger, 2016, Henriques & Moran, 2007, McKenney *et al.*, 2013, Stewart, 2015). The coat is differentiated into an inner, lamellar layer, and a more external, electrondense outer layer. The crust is formed by several cysteine-rich proteins that self-assemble into a hexagonal lattice, is glycosylated and stands in close apposition to the underlying outer coat (Jiang *et al.*, 2015, McKenney *et al.*, 2010, Waller *et al.*, 2004). In other sporeformers, the coat is surrounded by an exosporium, best characterized in spores of the pathogens *B. anthracis* and *B. cereus*, where it consists of a loose balloon-like structure separated from the edge of the coat by an interspace (reviewed by (Stewart, 2015)). The exosporium has a basal layer, which contains cysteine-rich proteins related to the crust proteins of *B. subtilis*, and that also assemble into a porous hexagonal lattice; a hair-like protrudes from the basal layer (Jiang *et al.*, 2015, Kailas *et al.*, 2011); reviewed by (Stewart, 2015)). The hairy nap is formed mainly by radial projections of a long collagen-like glycosylated protein, BclA, which is anchored in the basal layer (Stewart, 2015). While the exosporium contributes to the exclusion of lytic enzymes and antibodies, BclA is involved in a specific interaction with receptors at the surface of professional macrophages, essential for the course of infection in *B. anthracis* (Stewart, 2015). An exosporium that fits the general structural organization found in the *B. anthracis/B. cereus* group of organisms is present in several species of *Bacillus* and *Clostridium* (Stewart, 2015).

The composition, assembly and structure of the surface of *C. difficile* spores is of interest, firstly because infection begins with the ingestion of spores that have to resist the gastric barrier and secondly, because germination, which takes place in the intestine in response to bile salts (Sorg & Sonenshein, 2008) (Fig. 1B) and that may occur in close contact with host cells (Panessa-Warren *et al.*, 1997), is influenced by the structure and composition of the coat and exosporium (Barra-Carrasco *et al.*, 2013). Moreover, spores bind to colonic cells and to components of the intestinal mucosa and binding is influenced by the status of the surface layers (Barra-Carrasco *et al.*, 2013, Joshi *et al.*, 2012, Panessa-Warren *et al.*, 1997); binding to cells may involve a specific spore receptor as it reaches saturation (Mora-Uribe *et al.*, 2016) (Fig. 1B). Finally, the interaction of spores with host cells, mediated by the spore surface layers, is important for colonization and infection (Calderon-Romero *et al.*, 2018, Hong *et al.*, 2017, Phetcharaburanin *et al.*, 2014). *C. difficile* spores have a lamellar coat adjacent to the cortex, and an external, more electrondense layer surrounded by an exosporium closely apposed to the underlying coat (Pereira *et al.*, 2013, Phetcharaburanin *et al.*, 2014, Pizarro-Guajardo *et al.*, 2016a, Rabi *et al.*, 2017). At least some strains produce, in the same culture, two main spore morphotypes, with either a thick or a thin exosporium (Pizarro-Guajardo *et al.*, 2016a). How these morphotypes are produced, and the significance of this differentiation at the population level is unknown. Moreover, in spores produced by most clinical isolates, the exosporium is thicker than in a standard laboratory strain such as 630Δ*erm*, has a convoluted or bumpy appearance, and a hairy-like nap protrudes from its surface (Joshi *et al.*, 2012, Pizarro-Guajardo *et al.*, 2016a, Rabi *et al.*, 2017). *C. difficile* possesses three homologues of BclA, which form the hair-like nap of the exosporium (Phetcharaburanin *et al.*, 2014, Pizarro-Guajardo *et al.*, 2014).

At least BclA1 is required for colonization of the mouse gastro-intestinal tract, suggesting that the protein and/or proper assembly of the exosporium layer is important during the initial stages of infection (Phetcharaburanin *et al.*, 2014). Other exosporium proteins have been identified and at least two of these proteins, CdeC and CdeM, are cysteine-rich (Barra-Carrasco *et al.*, 2013, Calderon-Romero *et al.*, 2018, Diaz-Gonzalez *et al.*, 2015). Importantly, a *cdeC* mutant forms spores that lack the exosporium and the hairy nap, are more permeable to lysozyme and solvents and show altered binding to epithelial cells and to components of the intestinal mucosa (Barra-Carrasco *et al.*, 2013, Mora-Uribe *et al.*, 2016). *cdeM*, in turn, was previously shown to be strongly induced *in vivo*, upon infection of axenic mice with vegetative cells of *C. difficile* and a *cdeM* insertional mutant is impaired in colonization (Janoir *et al.*, 2014). CdeM is an important determinant of exosporium assembly, and also modulates the ability of spores to adhere to the mouse colonic mucosa (Calderon-Romero *et al.*, 2018).

Here we show that CdeM is a critical determinant of the assembly of a spore polar appendage that is continuous with and an integral part of the exosporium. In a *cdeM* mutant, the exosporium is thin and when present, the polar appendage is short and disorganized. Consistent with a role in the assembly of the exosporium and the polar appendage, we show that CdeM localizes around the spore and at the appendage, and contributes to the rigidity of spores. We found that in both the wild-type strain of a *cdeM* mutant, the spore population consists of spores with a well developed appendage or without an appendage or a much reduced one. *cdeM* spores with a short appendage germinate better that those without an appendage, but the two types of spores show enhanced germination compared to their wild-type counterparts, pointing to a direct role of the appendage in germination. While misassembly of the spore surface layers may explain the previously described colonization defect of a *cdeM* insertional mutant in a mouse axenic model (Janoir *et al.*, 2014), the mutant shows increased virulence in a hamster model of disease presumably because increased spore permeability results in enhanced germination. We suggest that the two types of spores correspond to distinct modes of germination. Since the representation of the two main spore types differs in an epidemic strain, their proportion in a population maybe an important trait for colonization and virulence.

## Results

### *Transcription of* cdeM *is directly controlled by* σ^K^

The proteins that compose the spore surface structures are produced in the mother cell, under the control of σ^E^ and σ^K^ in both *B. subtilis* (Henriques & Moran, 2007, McKenney *et al.*, 2013) and *C. difficile* (Fimlaid *et al.*, 2013, Pereira *et al.*, 2013, Pishdadian *et al.*, 2015, Putnam *et al.*, 2013, Saujet *et al.*, 2013). The *cdeM* gene, was proposed to be controlled by σ^K^ as *cdeM* expression is severely curtailed in a *sigK* mutant (Fimlaid *et al.*, 2013, Saujet *et al.*, 2013). To more precisely define the requirements for *cdeM* expression, we mapped its promoter using 5′-RACE. We identified a single transcriptional start site, a T, located 45 nt upstream of the predicted start codon of *cdeM*. We found upstream of this nucleotide at position -10 and -35 sequences that conform to the consensus elements of promoters recognized by σ^K^ in *C. difficile* (Fig. 2A) (Fimlaid *et al.*, 2013, Saujet *et al.*, 2013). In addition, we used quantitative RT-PCR (qRT-PCR) to monitor expression of *cdeM* during sporulation of a wild type (WT) strain (630Δ*erm*) or *sigF*, *sigE*, *sigG*, and *sigK* mutants. We have previously shown that S medium (SM; (Wilson *et al.*, 1982)) supports sporulation and that the activity of σ^F^, σ^E^, σ^G^ and σ^K^ can be detected and quantified between 14 h and 24 h of growth in this medium (Pereira *et al.*, 2013, Saujet *et al.*, 2013). We found that expression of *cdeM* was strongly reduced both in a *sigK* mutant (100-fold in transcriptome and 410-fold by qRT-PCR; Fig. 2B), unable to produce σ^K^, and in a *sigE* mutant (100-fold in transcriptome and 500-fold by qRT-PCR), which is required for the production and activity of σ^K^ (Fimlaid *et al.*, 2013, Pereira *et al.*, 2013, Pishdadian *et al.*, 2015, Serrano *et al.*, 2016). These results are in line with the presence of a promoter recognized by σ^K^ upstream of *cdeM.* In addition, expression of *cdeM* was not affected in a mutant unable to produce σ^G^, the late forespore-specific sigma factor (Fig. 2B) which, unlike in *B. subtilis*, is not required for the activation of σ^K^ in the mother cell in *C. difficile* (Fimlaid *et al.*, 2013, Pereira *et al.*, 2013, Pishdadian *et al.*, 2015, Saujet *et al.*, 2013, Serrano *et al.*, 2016). *cdeM* expression was also reduced in a *sigF* mutant (30-fold in transcriptome and by qRT-PCR) but to a lower extend compared to the effect of σ^E^ and σ^K^ (Fig. 2B). A control exerted by *sigF* over σ^K^ was also observed for other σ^K^-target genes (Saujet et al, 2013); it may result from either a partial control of σ^E^ activity by σ^F^ ((Fimlaid *et al.*, 2013, Saujet *et al.*, 2013) or to a still uncharacterized mechanism of control of σ^K^ activity by σ^F^. Together, these results indicate that σ^K^ directly transcribes the *cdeM* gene.

**Figure 2.**
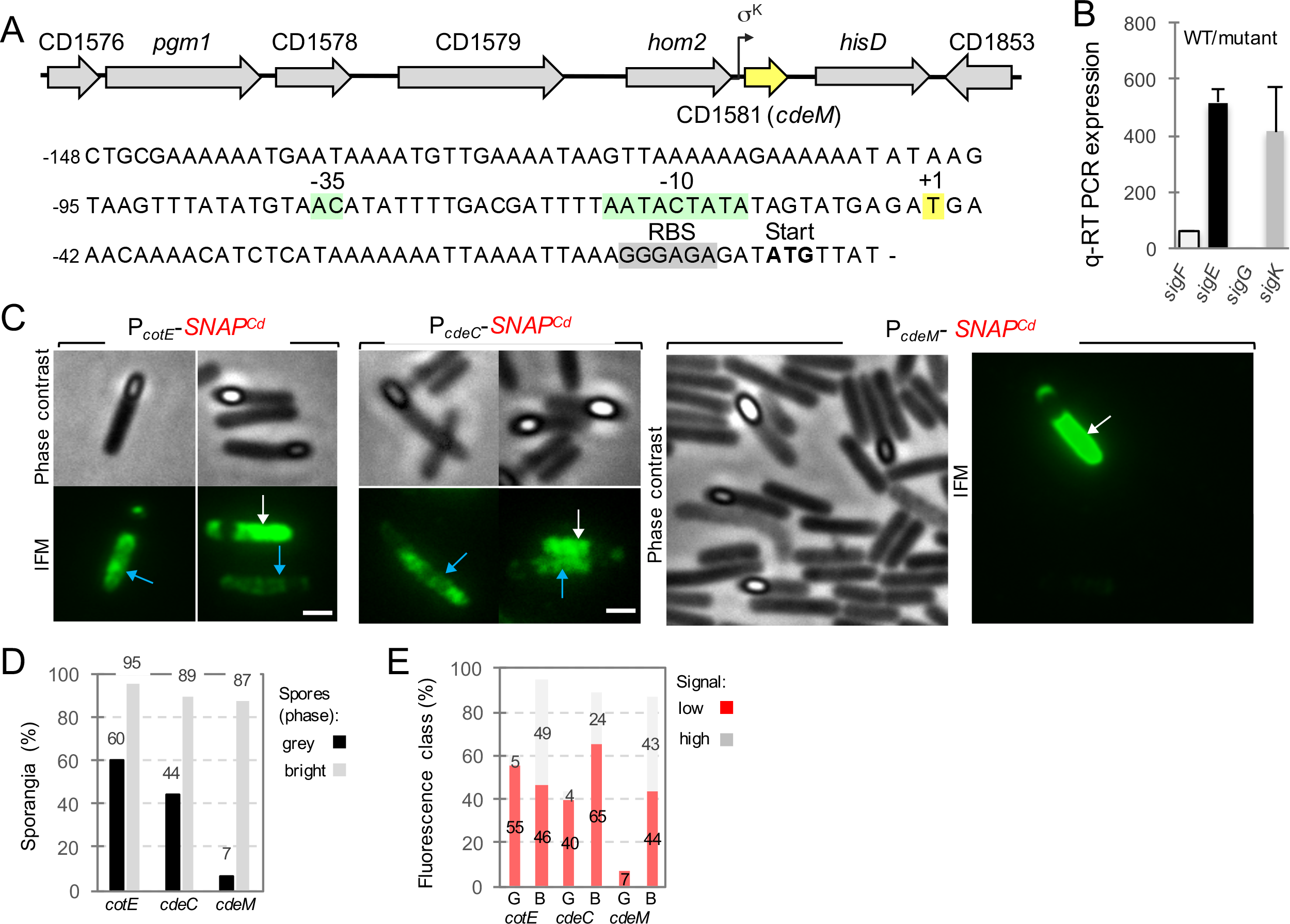
*cdeM* is expressed in the mother cell during late stages of spore morphogenesis. **A**: the *cdeM* region of the chromosome of strain 630Δ*erm*, encompassing 11 kb of DNA, and the *cdeM* promoter region. The transcription start site (+1, yellow), as mapped by RNAseq, and the -10 and -35 promoter elements (green) that match the consensus for σ^K^ binding are indicated. The ribosome binding site (RBS, shaded in grey) and the *cdeM* start codon are also indicated. **B**: Effect of mutations on the indicated regulatory genes on the level of the *cdeM* transcript as measured by qRT-PCR and represented as the WT/mutant ratio. **C**: expression of transcriptional P_*cotE*_-*SNAP*^*Cd*^, P_*cdeC*_-*SNAP*^*Cd*^, and P_*cdeM*_-*SNAP*^*Cd*^ fusions in relation to spore morphogenesis. The cells were collected after 24h of growth in SM liquid medium and subject to immunofluorescence microscopy (IMF) with an anti-SNAP antibody. Arrows: blue, phase grey spore; red, phase bright spore; white, strong mother cell-specific signal; grey, weak mother cell signal. Scale bar, 1 μm. **D**: the percentage of sporangia of phase grey and phase bright spores expressing P_*cotE*_-*SNAP*^*Cd*^, P_*cdeC*_-*SNAP*^*Cd*^, or P_*cdeM*_-*SNAP*^*Cd*^is shown. **E**: the intensity of the fluorescence signal (low versus high; see material and methods for a definition) is shown for the sporangia of phase grey (G) or phase bright (B) spores, for strains expressing P_*cotE*_-*SNAP*^*Cd*^, P_*cdeC*_-*SNAP*^*Cd*^, and P*_cdeM_-SNAP*^*Cd*^ fusions, as indicated.

### cdeM *is expressed during the final stages in spore morphogenesis*

To determine the time of *cdeM* expression in relation to morphogenesis and to compare its expression with that of genes encoding other coat/exosporium proteins, we examined the expression of *cotE*, *cdeC*, and *cdeM* during sporulation using strains harboring P_*cotE*_-, P_*cdeC*_-, and P_*cdeM*_-*SNAP*^*Cd*^ transcriptional fusions ((Pereira *et al.*, 2013); this work). *SNAP*^Cd^ production was monitored by immunofluorescence microscopy using an anti-SNAP antibody. Under these conditions, we were only able to score sporangia following engulfment completion, in which phase dark or phase bright spores were visible (Fig. 2C). For all three genes, sporangia with weak and stronger fluorescence signals, always confined to the mother cell, were detected, and we defined two levels of fluorescence signal intensity [weak, lower than 1000 arbitrary units (AU) and high, equal or higher than 1000 AU; see also the Material and Methods section]. These two classes were scored independently as a function of the stage in morphogenesis (Fig. 2C; weak signal, blue arrows; white arrows, stronger signal). CotE was detected in 60% of the sporangia of phase grey spores of which 55% showed a weak fluorescence signal and only 5% showed a strong signal (the remaining showing no labeling) (Fig. 2D and E). CotE was also detected in 95% of the sporangia of phase bright spores, and 49% of those showed a strong signal, while 46% showed a weak signal (Fig. 2D and E). *cotE* is thought to be under the control of σ^k^ (Pishdadian *et al.*, 2015, Saujet *et al.*, 2013). Expression of P_*cotE*_-*SNAP*^*Cd*^, however, is detected prior to engulfment completion in some sporangia but increased following completion of engulfment when the spore turns phase grey, then phase white and finally phase bright (Fig. 2C-E). The early expression of P_*cotE*_-*SNAP*^*Cd*^ may result from utilization of the *cotE* promoter by σ^E^; alternatively, it is caused by low levels of σ^K^ activity in sporangia prior to engulfment completion (Pereira *et al.*, 2013, Saujet *et al.*, 2013, Serrano *et al.*, 2016).

*cdeC* is also under the control of σ^K^ but additionally requires the ancillary transcription factor SpoIIID (Pishdadian *et al.*, 2015, Saujet *et al.*, 2013). Expression of *cdeC* was only detected following engulfment completion (Fig. S1B). Expression was detected in 44% of the sporangia of phase grey spores (40% of which showed a weak signal and 4% a stronger signal, the remaining with no labeling) and in 89% of the sporangia of phase bright spores (of which 65 % showed a weak signal, 24% a strong signal, and the remaining no labeling) (Fig. 2D and E). Thus, the onset of *cdeC* expression coincides with the appearance of phase grey spores. Finally, expression of *cdeM* was also only detected following engulfment completion (Fig. 2C-E) in agreement with the presence of a σ^K^-dependent promoter (Fig. 2A). Expression of *cdeM* was detected in 7% of the sporangia of phase grey spores (all of which showed a weak signal) but increased to 87% of the sporangia of phase bright spores (of which 44% showed a weak signal and 43% a strong fluorescence signal) (Fig. 2D and E). Although the percentage of sporangia showing a fluorescence signal was lower, the analysis of P_*cotE*_-*SNAP*^*Cd*^, and P_*cdeM*_-*SNAP*^*Cd*^ and P_*cdeC*_-*SNAP*^*Cd*^ expression using the TMR-Star SNAP substrate confirmed the results obtained using immunofluorescence (Fig. S1).

While the transcription of genes such as *cotE* begins prior to engulfment completion, *cdeC* is expressed after engulfment completion. *cdeM* in turn, is activated only at late stages in spore morphogenesis, when the spore becomes phase bright, just prior to lysis of the mother cell. Furthermore, the data also suggests that at least following engulfment completion, the levels of expression of *cotE*, *cdeC* and *cdeM* show some heterogeneity across the cell population.

### CdeM is an abundant component of the spore surface layers that undergoes multimerization

As a first step to characterize the role of *cdeM* we characterized spores of a previously constructed insertional mutant, *cdeM::ermB*, constructed in the background of strain 630Δ*erm* (Janoir *et al.*, 2014). We first examined the proteins extractable from mature spores under our experimental conditions. Spores were purified to greater than 99% homogeneity by density centrifugation on gradients of metrizoic acid (see Materials and Methods). A collection of about 30 polypeptides was extracted from wild-type (WT) spores as revealed by Coomassie blue staining (Fig. 3A, lane 1). The pattern obtained differs slightly from other recent reports in the number of bands and their relative abundance. These differences likely reflect different culturing, spore purification and extraction conditions. Nevertheless, 11 proteins were identified by mass spectrometry analysis, which were previously found to be spore surface components or part of the spore proteome (Abhyankar *et al.*, 2013, Barra-Carrasco *et al.*, 2013, Calderon-Romero *et al.*, 2018, Diaz-Gonzalez *et al.*, 2015, Escobar-Cortes *et al.*, 2013, Lawley *et al.*, 2009, Paredes-Sabja & Sarker, 2012, Permpoonpattana *et al.*, 2011). These include Pfo (a pyruvate-ferredoxin oxidoreductase, coded for CD2682, below and above the 250 kDa marker; predicted molecular mass of 128 kDa), AdhE, a bifunctional acetaldehyde CoA/alcohol dehydrogenase (95 kDa, coded for by CD2966), CotE (coded for by CD1433; predicted size of 82 kDa, but detected in the 40 kDa region of the gel possibly due to proteolytical separation of the peroxiredoxin and chitinase domains; (Permpoonpattana *et al.*, 2013, Permpoonpattana *et al.*, 2011)), CdeC (42 kDa, coded for by CD1067; detected in both the 80 and 200 kDa regions of the gel), CotB (35 kDa, CD1511), CotA (34 kDa, CD1613), CdeM (at 17 kDa; predicted molecular weight of 19 kDa; coded for by CD1581), SleC (CD0551), a rubrerythrin (Rbr; 25 kDa/CD0825) and CotCB (21 kDa, CD0598) (Fig. 3A, in the WT, red arrows). CdeM, which was also detected as an abundant species in WT spores, was absent from extracts prepared from spores of the *cdeM*::*ermB* mutant (Fig. 3A, green arrows). Importantly, complementation of *cdeM::ermB* in single copy using transposon Tn*916* (this strain is henceforth referred to as *cdeM*^C^; see Fig. S2) or a multicopy plasmid (pFT43) restored the presence of the protein at the spore surface (Fig. 3A). The absence of CdeM was the main difference between the protein profile obtained for the WT and *cdeM*::*ermB* spores (Fig. 3A). However, the extractability of at least two other species, CotJC1 and SleC, increased in the *cdeM*::*ermB* mutant, while the extractability of several others, including NifJ and CotB, decreased (Fig. 3A). Reduced extractability of CotB in *cdeM* spores was also reported in a recent study (Calderon-Romero *et al.*, 2018). These authors also reported a significant decrease in the levels of two forms of CdeC (of 140 and 44 kDa), in spores of the *cdeM* mutant (Calderon-Romero *et al.*, 2018). In our hands, CdeC was detected as a species of about 52 kDa, which most likely corresponds to the 44 kDa species detected by Calderón-Romero and co-authors (the predicted molecular weight of CdeC is 44.7 kDa). The extractability remained unchanged, relative to the WT, in *cdeM*::*ermB* spores (Fig. 3B, bottom panel).

**Figure 3.**
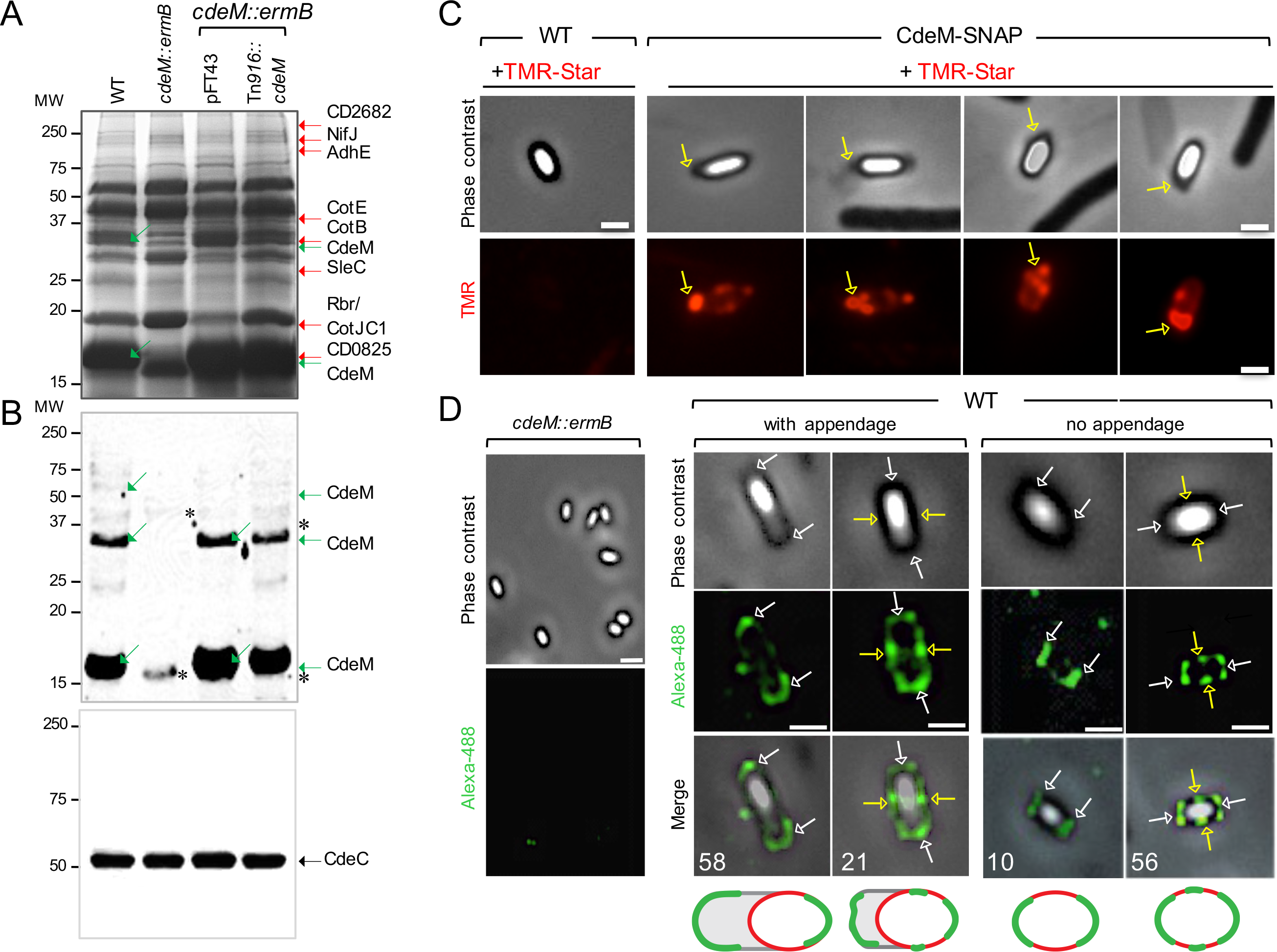
Localization of CdeM in spores. **A**: shows the SDS-PAGE analysis of the proteins extracted from purified spores of the wild type strain 630Δ*erm* (WT), the *cdeM::ermB* mutant, and the mutant complemented with either *cdeM* in a multicopy plasmid (pFT43) or integrated into the genome in single copy (Tn*916::cdeM*). The gel was stained with Coomassie. Red arrows indicate proteins, identified by mass spectrometry in the WT (see text for details); the green arrows indicate the position of forms of CdeM. **B**: Immunoblot analysis of the gel in A with anti-CdeM (top) or anti-CdeC (bottom) antibodies. The black arrows indicate CdeC; in A and B, the green arrows indicate forms of the CdeM protein and the asterisks indicate species that cross-react with the anti-CdeM antibody. The position of molecular weight (MW) markers (in kDa) is indicated. **C**: localization of CdeM-SNAP^Cd^ in spores by fluorescence microscopy. Spores of a 630Δ*erm* derivative producing CdeM-SNAP were labeled (+TMR-Star) or not (no TMR-Star) with the SNAP substrate TMR-Star SNAP and examined by fluorescence microscopy. The yellow arrows point to the strong signal associated with one of the spore poles. **D**: immunofluorescence analysis of WT and *cdeM*::*ermB* spores using an anti-CdeM antibody and an Alexa-488-conjugated secondary antibody. The yellow arrows point to a polar florescence signal and the white arrows to signals located along the long axis of the spore. The numbers refer to the percentage of spores showing the pattern represented in schematic form below. Scale bar in C and D, 1 μm.

In addition to the 17 kDa, possibly monomeric form of CdeM, the immunoblot analysis of the WT spore extracts using a polyclonal antibody raised against the partially purified protein (see the Material and Methods), revealed additional forms of about 25 kDa, 37 kDa and 60 kDa (Fig. 3B). CdeM contains 14 cysteine residues, some of which part of repeat motifs previously noticed (Calderon-Romero *et al.*, 2018); 7 of the cysteine residues are clustered close to its C-terminal end, from residues 151 through 160 of the 161 residues-long protein (Fig. S3A and B). The Cys-rich C-terminal end of CdeM is preceded by a polymorphic region (Fig. S3B, region P4; see also the supplemental text). The partially purified His_6_-CdeM protein runs at about 17 kDa in SDS-PAGE in the presence of reducing agents (Fig. S3C). In the absence of reducing agents, however, CdeM runs as a major species at about 60 kDa, as well as several other species of even higher apparent molecular weight (Fig. S3C). Therefore, the 25 kDa, 37 kDa and 60 kDa forms detected in the spore extracts by immunoblotting are likely to be disulfide-cross-linked homo-multimeric forms of CdeM. These species are at least partially resistant to boiling and the reducing SDS-PAGE conditions.

### CdeM localizes asymmetrically at the spore surface

We next wanted to localize CdeM at the surface of mature spores. We first used a CdeM-SNAP^Cd^ translational fusion produced from a low copy plasmid in an otherwise WT background. Labeling of the purified spores with the TMR-Star substrate reveals the preferential localization of CdeM-SNAP^Cd^ at one or both of the spore poles (Fig. 3C, yellow arrows). Often, the signal overlapped with an appendage, visible in the phase contrast images, protruding from one of the spore poles, suggesting the presence of CdeM-SNAP^Cd^ in this structure (Fig. 3C, yellow arrows; see also below). The SNAP-TMR signal is specific, since no fluorescence is detected in spores of the WT strain, which does not produce SNAP^Cd^, labeled with TMR-Star (Fig. 3C, left panels).

To localize the native protein, we used immunofluorescence microscopy. The polyclonal anti-CdeM antibody was bound to WT and *cdeM::ermB* mutant spores, and the antigen-antibody complex detected with a secondary anti-rabbit fluorescent antibody. We found two main types of spores in the population, those with a visible polar appendage and those without (see also the following section for a quantitative description of this phenotype). Both types of spores were decorated by the antibody. In 79% of the spores with a polar appendage, the antibody decorated the tip of the appendage, but not at the appendage body, and also the opposite spore pole (Fig. 3D, white arrows). Among the spores with an appendage, 21% showed a somewhat shorter appendage, and in those spores CdeM, was also found at the spore body (Fig. 3D, yellow arrows). In 10% of the spores without a visible appendage, CdeM localized at the two spore poles, whereas in 56% of those spores CdeM showed a punctate pattern of fluorescence around the entire contour of the spore, with the remaining showing no decoration (Fig. 3D). Importantly, only residual unspecific labeling of *cdeM*::*ermB* spores was detected (Fig. 3D, right). The decoration of WT spores by the anti-CdeM antibody, along the periphery of the spores and at the edge of the appendage is consistent with the proposal that CdeM is at least partially exposed at the spore surface, and accessible to the antibody (Barra-Carrasco *et al.*, 2013, Calderon-Romero *et al.*, 2018). Consistent with this inference, incubation of purified WT or TMR-labelled CdeM-SNAP^Cd^ spores with trypsin reduced the amount of extracted CdeM (Fig. S4A) and the intensity of the fluorescence signal (Fig. S4B). CdeM however, is tightly associated with the spore surface, as it was not significantly released from spores by washing with a 1M solution of potassium chloride (Fig. S4C). Also consistent with a surface localization, *cdeM*::*erm* spores showed reduced hydrophobicity (Fig. S4D). Importantly, the comparison between labeling of CdeM-SNAP^Cd^ with the small molecule TMR, and the decoration pattern obtained by immunofluorescence suggests that CdeM localizes both at the edge and the body of the appendage, but is not accessible or not recognized by the antibody at the body of this structure. In any case, CdeM is an abundant component of the spore surface (Barra-Carrasco *et al.*, 2013, Calderon-Romero *et al.*, 2018) that undergoes multimerization at the spore surface, and that localizes both at the polar appendage as well as around the spore body.

### CdeM is a key determinant of assembly of a spore polar appendage

The observation that a population of WT spores consisted of a mixture of appendage-bearing and appendage-lacking spores (above), together with the polar localization of CdeM suggested that the protein could have a role in the assembly of this structure. We have shown before that the lipophilic dye FM4-64 stains spores of *C. difficile* (Pereira *et al.*, 2013) and we used this dye in combination with phase contrast microscopy to characterize populations of WT and *cdeM*::*ermB* mutant spores. In both WT and *cdeM* spores, the polar appendage is clearly distinguishable from the spore body, as it does not stain with FM4-64 (Fig. 4A). The most prevalent appendage pattern seen for the WT is a relatively long, well-organized and partially refractile appendage (Fig. 4A, type A: about 25% of the population) whereas only 3% of the spores showed a short, disorganized appendage (type B), with the remaining spores (72% of the population) showing no appendage (Fig. 4A). In contrast, type B appendages were the most prevalent for *cdeM*::*ermB* spores (about 20%) whereas only 7% showed a type A appendage, while the remaining spores (73% of the population) showed no appendage (Fig. 4A; see also the Supplemental text). The WT numbers were restored in the *cdeM*^C^ strain (Fig. 4A and B). Interestingly, the percentage of type A spores was higher for the epidemic strain R20191 (Fig. S5A; see also the supplemental text).

**Figure 4.**
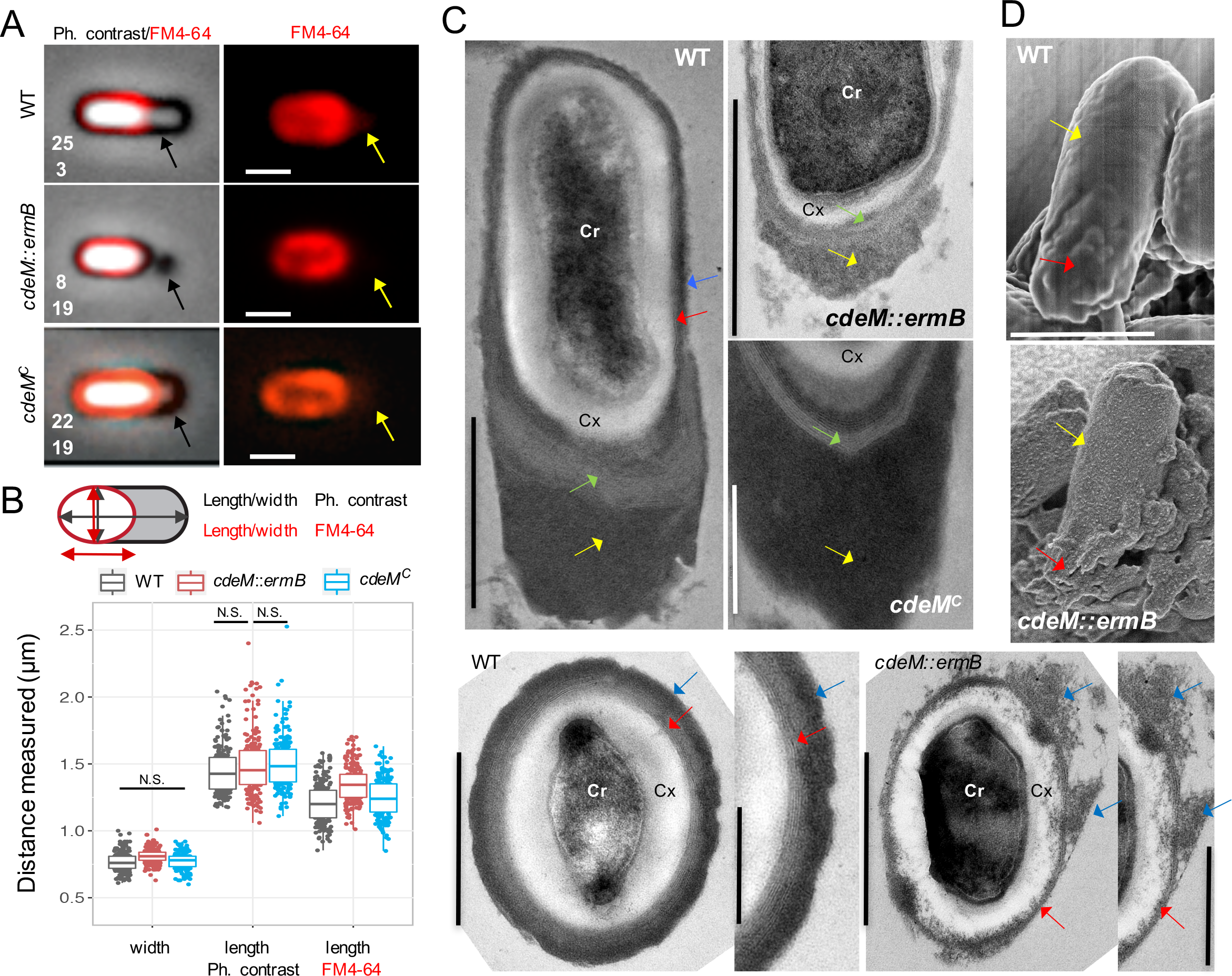
CdeM is required for proper assembly of a spore polar structure. **A**: Phase contrast (PC) and fluorescence microscopy images of the two main types of spores produced by a WT strain, the *cdeM* mutant and the complementation strain (Tn*976*::*cdeM*, or *cdeM*^C^) following staining by FM4-64. The black arrow points to a polar structure that in the WT and the complementation strain is partially refractile (type A appendage); this structure is shorter and phase dark in the *cdeM*::*ermB* mutant (type B appendage). Note that neither type of appendages stains with FM4-64 (yellow arrow). The numbers refer to the percentage of type A (top) and type B (bottom) appendages scored for each strain. Scale bar, 1 μm. **B**: The top diagram shows the measures of length and width measured for spores in phase contrast (black) and FM4-64 images (red). Note that the width coincides for the images. The bottom panel is a box plot showing the median, lower and upper quartiles of the distance measured (in μm), for each of the indicated strains, in the phase contrast or FM4-64 images. Each point is a measurement of a single spore. Pairwise differences between the strains are statistically significant for all measures (*P* < 0.05; Kruskal-Wallis rank sum test for width and length Ph. Contrast; ANOVA for length FM4-64), except for those comparisons denoted by a black bar connecting them and labeled by N.S. (not significant). **C**: transmission electron microscopy of thin sections of WT, *cdeM*::*ermB* and *cdeM*^C^ spores. The two top rows of panels show TEM images of longitudinal sections whereas the third row are cross sections. The yellow arrows point to the polar appendage region and the green arrow points to a region between the coat and the appendage seen in WT spores; the red arrows point to the lamellar coat, and the blue arrow to the more external, exosporium-like layer, that is reduced in *cdeM*::*ermB* spores, or to the electrondense material that remains attached to *cdeM* spores. Cr, spore core; Cx, spore cortex. Scale bar for the three top rows of panels, 0.2 μm. The two bottom panels are SEM images of WT and *cdeM* spores. The yellow arrow points to the spore body and the red arrow to the appendage. Scale bar for the two bottom panels, 1.0 μm. D: scanning electron microscopy of a WT and a *cdeM*::*erm* spore. The yellow label points to the spore body and the yellow arrow to the appendage region. Scale bar, 0.2 μm.

As a further characterization of the appendage, we measured the spore length in both the phase contrast and FM4-64 images for individual spores as well as the spore width (Fig. 4B, top). The length measurement in the phase contrast images gives the total length of the spore (spore body and appendage) whereas the FM4-64 images gives the length of the spore body (without the appendage), regardless of the appearance of the appendage (Fig. 4B, top). The spore width measurement coincides in the two images (Fig. 4B, top). The spore length, as measured in the phase contrast images did not differ much among spores of the three strains (Fig. 4B; WT: 1.45±0.17 μm, *cdeM*: 1.49±0.22 μm; *cdeM*^C^: 1.51±019 μm). This is consistent with the presence of an appendage, albeit disorganized, in *cdeM* spores (Fig. 4A) and suggests that *cdeM* affects mainly the organization of this structure, and not its presence or absence. In contrast, the length and width of the spore body, measured in the FM4-64 images, is larger for *cdeM* spores than for WT or *cdeM*^C^ spores (Fig. 4B; L: 1.35±0.13 μm/W:0.81±0.06 μm for *cdeM*; L: 1.20±0.15 μm/W:0.77±0.07 μm for WT; L: 1.25±0.14 μm/W:0.77±0.06 μm for *cdeM*^C^). This suggests that expression of *cdeM* imposes a constraint on the dimensions of the spore and that in the absence of CdeM some expansion of the spore body occurs. We also note that spores of strain R20291, which have a thicker exosporium layer as compared to 630Δ*erm* spores (Pizarro-Guajardo *et al.*, 2016a, Pizarro-Guajardo *et al.*, 2014, Pizarro-Guajardo *et al.*, 2016b), also have a longer appendage (Fig. S5B; see also the supplemental text).

To next used transmission electron microscopy (TEM) to examine the impact of the *cdeM* insertional allele on the structure of the spore. The coat of *C. difficile* spores shows an inner and an outer layer, both lamellar, with the outer coat more electrondense; an electrodense exosporial layer surrounds the spore, closely apposed to the coat (Lawley *et al.*, 2009, Paredes-Sabja & Sarker, 2012, Pereira *et al.*, 2013, Permpoonpattana *et al.*, 2011, Rabi *et al.*, 2017). For WT spores, a lamellar coat layer in close contact with the spore cortex peptidoglycan (Fig. 4C, red arrows) and the electrodense exosporial layer attached to the coat were seen (Fig. 4C, blue arrows). The distinction between an inner and an outer coat layer, however, was not evident (Fig. 4C). In longitudinal sections of WT spores, a compact polar appendage is clearly seen (Fig. 4C, yellow arrow); a basal region at the spore/appendage junction is also discernible (green arrow). Importantly, the TEM images clearly show that the appendage is continuous with the exosporium. In *cdeM* spores the appendage appears shorter and less electrondense in *cdeM* spores and the spore/appendage junction region is poorly defined (Fig. 4C). In cross sections, the coat and the exosporium appear well defined for WT spores (Fig. 4C, bottom panels). In contrast, in *cdeM* spores, the coat layer appears thinner, and the exosporium is largely absent (Fig. 4C). These two features, a thin coat and the absence of the exosporium have also been described by Calderón-Romero and co-workers (Calderon-Romero *et al.*, 2018). The outer layer of the coat, however, which these authors reported as being thicker in the *cdeM* mutant, was not evident (Fig. 4C), suggesting that under our experimental conditions (including sample preparation), this structure is either not formed or not retained by spores when most of the exosporium is absent. Nevertheless, disorganized material, most likely remnants of the outer coat layer and/or exosporium, remain loosely attached to the exposed edge of the inner coat (Fig. 4C). The structural features seen in the spore cross sections are in agreement with previous descriptions of WT spores (Barra-Carrasco *et al.*, 2013, Calderon-Romero *et al.*, 2018, Diaz-Gonzalez *et al.*, 2015, Escobar-Cortes *et al.*, 2013, Pereira *et al.*, 2013, Pizarro-Guajardo *et al.*, 2016a, Pizarro-Guajardo *et al.*, 2016b). Polar spore appendages have been described in other organisms, including *C. bifermentans*, *C. sporogenes* and *C. sordellii* (Panessa-Warren *et al.*, 1997, Rabi *et al.*, 2017, Samsonoff *et al.*, 1971). A polar appendage has been suggested to form as the result of a reorganization of the spore surface during germination of *C. difficile* spores (Panessa-Warren *et al.*, 1997), but has not been described, to our knowledge, as part of mature *C. difficile* spores. This structure is also clearly seen in scanning electron microscopy (SEM) images of WT and *cdeM* spores, although it is disorganized in the latter case (Fig. 4D, red arrows in the two bottom panels).

In any event, the requirement for CdeM for proper assembly of the exosporium and the polar appendage, is in line with the dual localization pattern of CdeM or CdeM-SNAP^Cd^ along the spore body and at the appendage region (Fig. 3C and 3D). Because CdeM is the most abundant protein extracted from WT spores that is absent from *cdeM*::*erm* spore extracts, we infer that the protein is likely to be the main structural component of the exosporium, which comprises the polar appendage.

### CdeM contributes to the rigidity of spores

Assembly of the exosporium and appendage may afford rigidity to the spore, as suggested by the increased length and width of the spore body in *cdeM* spores (above). To test this possibility, we used atomic force microscopy (AFM) to image purified WT and *cdeM*::*erm* spores. Topography (Fig. 5A, top panel), amplitude (middle) and phase (bottom) images show a relatively smooth surface in WT spores with a well-organized rigid polar region (Fig. 5A, white arrows; inserts show the spore polar region at a higher magnification). The surface of *cdeM* spores was more irregular, with a less organized and softer polar region from which material seems to peel-off possibly through the action of the microscope tip (Fig. 5A, white arrows in the amplitude images). The phase images also reveal a region of decreased rigidity that runs along the longitudinal axis of WT spores (Fig. 5A, white arrows in the phase panel). Imaging of this region at higher magnification suggests the presence of individual protein complexes along the two sides of a crevice (Fig. 5A; black arrows in the two lower panels). Importantly, this feature could not be defined for the *cdeM* mutant, as the entire spore surface, including the polar region, is softer (Fig. 5A). In all, the AFM results are in line with the view that CdeM participates in the assembly of the entire surface of the spore, appendage included, and that it confers rigidity to the spore.

**Figure 5.**
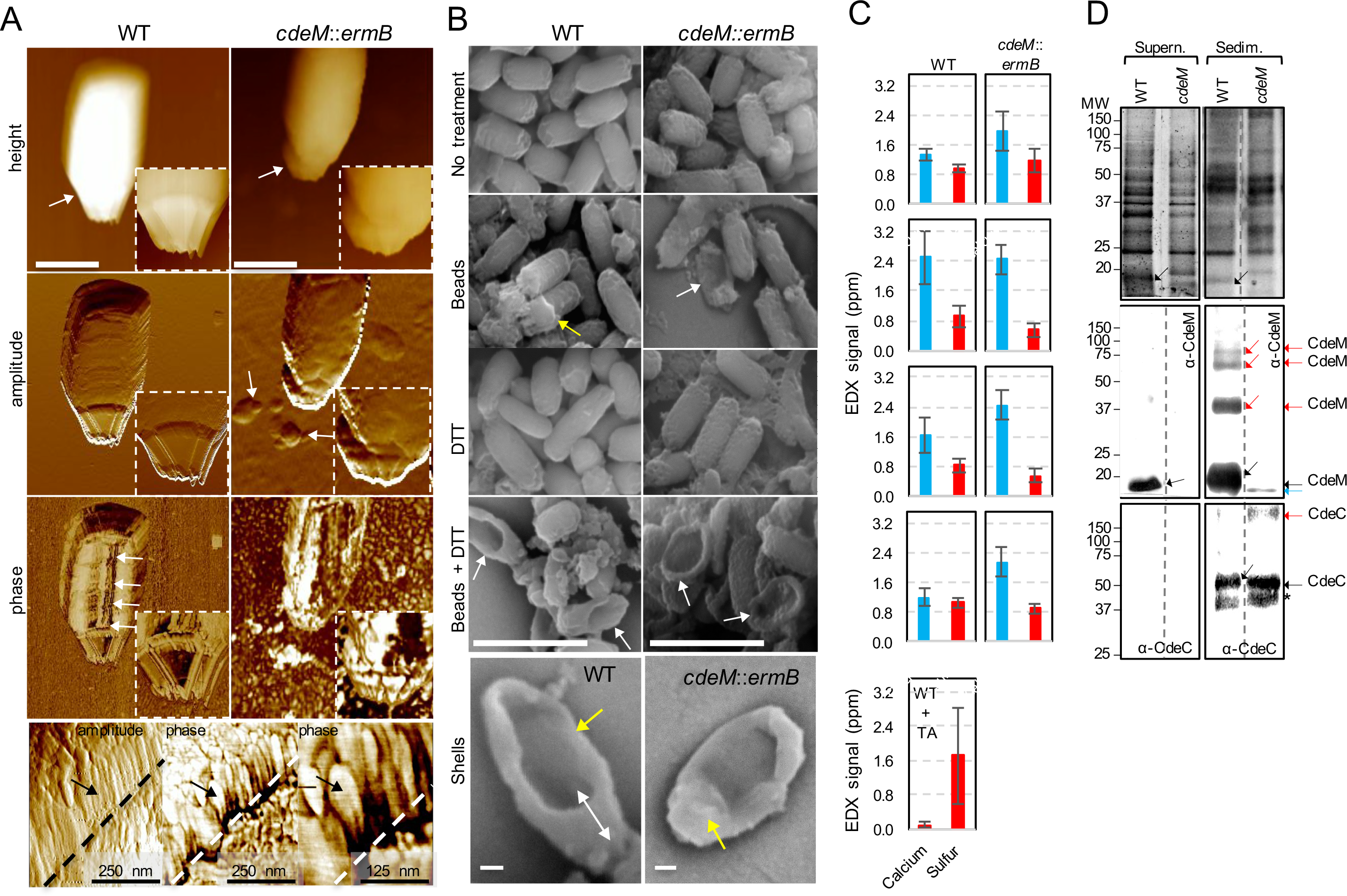
Surface features of WT and *cdeM*::*ermB* spores. **A**: atomic force imaging of purified WT and *cdeM*::*ermB* spores. Shown are topography (top panel), amplitude (middle) and phase (bottom) images. The amplitude image is a spatial derivative of the topography image, which is very sensitive to small, sharp features. The phase image depicts the phase shift between the excitation signal and the tip oscillation response, and it is sensitive to surface hardness. The white arrows in the amplitude images point to the spore polar appendage region. The white arrows in the phase image of WT spores point to a region of decreased rigidity that runs along the longitudinal axis of the spore. The inserts show the polar region of the spore. Scale bar, 1 μm. **B**: scanning electron microscopy analysis of purified WT and *cdeM::ermB* spores before (top row) or after treatment with beads (second row), 10 mM DTT (third row) and beads in the presence of 10 mM DTT (bottom row). The yellow arrow points to a spore with a partially disrupted shell; the white arrows point to empty shells. The two bottom panels show higher magnification images of the shells obtained following the beads/DTT treatment. Scale bar, 3 μm. **C**: shows the EDX analysis of the samples in B after treatment with beads in the presence of DTT. The EDX signal is shown for calcium (red) and sulfur (blue). The two bottom panels show the calcium and sulfur signals (same color code) after inducing germination of WT spores with 5% taurocholate (TA). **D**: suspensions of spores of the indicated strains were centrifuged and proteins in the supernatant or sediment fractions were extracted and resolved by SDS-PAGE (top panels; Coomassie-stained gels). The presence of the CdeM and CdeC proteins was then monitored by immunoblotting (middle and bottom panels). Black arrows show the position of CdeC or CdeM; red arrows show the position of multimeric species of CdeC or CdeM. The blue arrow shows the position of a species that cross-reacts with the anti-CdeM antibody and the asterisk the position of a possible CdeC degradation product. Molecular weight markers (in kDa) are shown on the left side of the panels.

### The spore appendage is an integral part of the exosporium

To test the hypothesis that the appendage is continuous with the exosporium, we attempted to isolate this structure. Earlier work has shown that sonication of spores in the presence of sarkosyl and proteases completely extracts the exosporium proteins, leaving behind a seemingly intact coat (Escobar-Cortes *et al.*, 2013). That several of the proteins assigned to the exosporium-like layer are Cys-rich, including CdeC and CdeM (Barra-Carrasco *et al.*, 2013, Diaz-Gonzalez *et al.*, 2015), together with the finding of a longitudinal region of lower rigidity in WT spores by AFM (above), suggested to us that the surface layer of spores could be sensitive to mechanical disruption in the presence of a reducing agent. Purified spores were shaken with either glass beads, the reducing agent DTT, or both (Fig. 5B). SEM of WT spores before any treatment revealed that most spores have a smooth surface, whereas the surface of *cdeM* spores has a porous appearance (Fig. 5B, top panels). Treatment of WT spores with glass beads results in the partial removal of an external layer, leaving behind an intact spore (Fig. 5B, second row, yellow arrow). In contrast, for *cdeM* spores, treatment with glass beads often resulted in the release of a complete empty shell; this shell shows a wide opening that appears to coincide with the position of the region of low rigidity seen for WT spores by AFM and that ends close to the junction between the polar appendage and the spore body (above) (Fig. 5B, second row, white arrow). This suggests that *cdeM* spores are less rigid or less stable than WT spores, in line with the light microscopy, TEM and AFM data (above). While treatment with DTT alone had no major impact of the surface of either WT or *cdeM* spores, the combined action of the beads and the reducing agent caused the release of empty shells in most of the spores of the two strains (Fig. 5B; white arrows in the fourth row of panels). We infer that the structural rigidity of spores is in part conferred by the formation of disulfide cross-links involving the Cys-rich exosporium proteins, among which CdeM.

Consistent with the view that CdeM is a key structural component of the exosporium, but not essential for its formation, the shells extracted from *cdeM* spores were thinner (0.063±0.008 μm) than for 630Δ*erm* spores; conversely, they were thicker for spores of epidemic strain R20291 (Fig. S5C and D; see also the supplemental text).

During none of the treatments did we see cells outgrowing from spores. Nevertheless, to test whether formation of the empty shells resulted from spore germination, we used SEM in combination with energy dispersed X-ray (EDX) spectroscopy. Release of Ca^2+^ from the spore core when germination is induced by the bile salt taurocholate (TA) results in a strong reduction of the signal for this element in spores, while the signal for sulfur remains essentially unchanged (Fig. 5C, bottom panels). Thus, we measured the Ca^2+^ and sulfur signals for WT and *cdeM* spores before and after treatment with glass beads in the presence of DTT. The sulfur signal, which is not expected to change during germination, showed variation among the different conditions tested but remained high following the different treatment; the signal for Ca^2+^ also showed some variation, but it never came down to nearly undetectable levels, as when germination is induced by TA (Fig. 5C, compare the three first sets of panels with the bottom panel, which shows measurements for WT spores following induction of spore germination with TA). Although we cannot exclude that some spores germinate under our experimental conditions, we infer that the empty shells result mainly from the mechanical/chemical detachment of the exosporium that surrounds the entire spore and is continuous with the polar appendage.

### *Evidence for two populations of CdeM* at the spore surface

Following treatment with the glass beads in the presence of DTT, the spore suspension was separated into a sediment containing spores and shells, and a supernatant, which we expected to contain proteins released from the spores during treatment. Several proteins were found in the supernatant, following SDS-PAGE and Coomassie staining, including, for WT spores, the 17 kDa form of CdeM; this species is also detected by immunoblotting and is absent from the supernatant of *cdeM* spores (Fig. 5D, black arrow in the top and middle panels). This species seemed reduced in the WT sediment by Coomassie staining relative to the supernatant, but reacted more strongly with the anti-CdeM antibody (Fig. 5D, top and middle panels). We do not presently know the reason for this apparent difference in recognition of CdeM in the supernatant and in the sediment by the antibody, but one possibility is that the protein is subject to a post-translational modification. That several multimeric forms of CdeM (close to the 37 kDa marker, as well as below and above the 75 kDa marker) are detected in the sediment from the WT (Fig. 5D, top and middle panels, red arrows) suggests that the shells are enriched in cross-linked forms of the protein. These multimeric forms of CdeM appear to be less extractable from whole spore preparations, *i.e.*, without any prior treatment (see Fig. 3A). CdeC was not detected in the supernatant of WT or *cdeM* spores, but was found as two species of about 60 kDa and 44 kDa in the sediment of both WT and *cdeM* spores and at slightly higher levels in the latter (Fig. 5D, black arrow in the bottom panel). The 44 kDa species likely corresponds to the low mass CdeC previously detected in spore extracts (Calderón-Romero et al., 2018). A high molecular weight form of CdeC, above the 150 kDa marker, was also detected in the *cdeM* spore sediment (Fig. 5D, red arrow in the lower panel). This form was not detected in the total spore extracts (Fig. 3A and B), suggesting that the shells are enriched in cross-linked forms of CdeC. The results suggest that CdeC is tightly associated with the spore and is not released into the supernatant by the glass beads/DTT treatment. Moreover, the increased extractability of CdeC from the shells isolated from *cdeM* spores suggests a more internal localization for CdeC and that CdeM, somehow reduces the extractability of CdeC. The CdeM protein, in contrast, exists as two populations; part is released to the supernatant upon treatment with beads/DTT, suggesting a looser association with the spore, while the spore and shell sediment is enriched in multimeric forms of the protein.

### Density gradient separation of different spore morphotypes

To determine whether the multimeric forms of CdeM were preferentially associated with spores bearing a polar appendage. We used density centrifugation to separate spores without an appendage from those bearing either short or long appendages. On 40%-60% step gradients of metrizoic acid, a WT population of spores forms bands at 55%, 50% and 45% (Fig. 6A, top). As determined by SEM, the initial material consisted of a heterogeneous population of spores while the denser fractions appeared enriched for appendage-bearing spores (Fig. 6A, bottom). In agreement with this, phase contrast microscopy shows that the spores collected from the band at 55% showed well developed appendages, as compared to those at 45% (Fig. S6). We measured the spore length in both the phase contrast and FM4-64 images as well as the spore width for individual spores in the initial material and in the various fractions, as described above (see also Fig. 4B). For both the WT and *cdeM* spores, the material at the 45% density layer consisted mainly of smooth spores that appeared to lack an external layer and have no appendage or a very short one, as assessed by SEM (Fig. 6A, top row of the SEM panels). For both the WT and *cdeM* spores, the width was similar to that in the initial material (Fig. 6B). The spore length, however, measured in the phase contrast images was reduced for both strains as compared with the initial material but increased in the FM4-64 images for *cdeM* spores (Fig. 6B; 1.13±0.18 vs 1.23±0.14 μm). This again suggests longitudinal expansion of the spore in the absence of CdeM (see also above). The 50% and 55% density fractions were enriched in spores that were longer than those in the 45% fraction as measured in the phase contrast images (Fig. 6B; e.g., length of WT spores: 1.10±0.15 μm in the 45% fraction vs. 1.35±0.18 μm in the 55% fraction). In line with the idea that CdeM imparts rigidity to the spore, *cdeM* spores were however shorter in the 50% and 55% fractions, as measured in the FM4-64 images (Fig. 6B; 1.23±0.14 μm in the 45% fraction vs. 1.06±0.11 μm in the 50% fraction). Together, this analysis suggests that spores in the 45% fraction may have a thin exosporium and lack an appendage, whereas spores in the 50% and 55% density fractions have a more developed exosporium and a polar appendage, even though disorganized in the case of the *cdeM* mutant.

**Figure 6.**
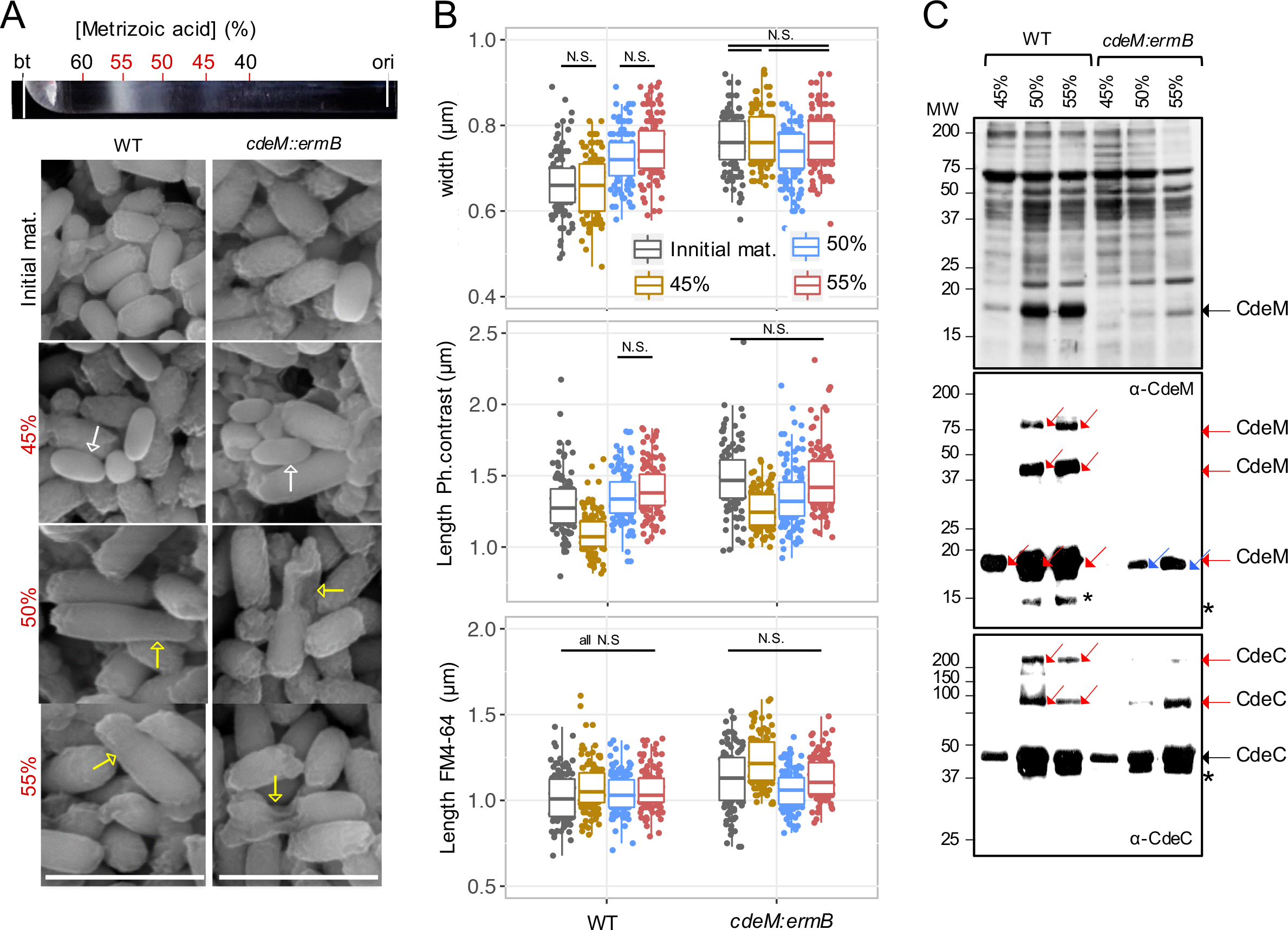
Appendage-bearing spores are enriched in multimeric forms of CdeM and CdeC. **A**: Purified spores of the WT and *cdeM*::*ermB* mutant were separated on discontinuous gradients of metrizoic acid (from 40 to 60%) and resolved in bands at 45%, 50% and 55% (the top panel illustrates the separation for WT spores). The bottom series of panels shows the analysis of the three fractions by scanning electron microscopy. The white arrow point to smooth spores, lacking an exosporium-like layer; the yellow arrows point to spores with an exosporium-like surface layer, which is continuous with the spore polar appendage. Scale bar, 2 μm. **B**: Box plots showing the median, lower and upper quartiles of the distance measured (in μm), for the initial spore suspension, and for the spores collected in the 45%, 50% and 55% fractions of the metrizoic acid gradients. Each point is a measurement of a single spore. Pairwise comparisons between fractions are statistically significant for both WT and *cdeM*::*ermB* spores (*P* < 0.05; Kruskal-Wallis rank sum test for length in the Ph.contrast and length in the FM4-64 images; ANOVA for width), except for those comparisons denoted by a black bar connecting them and labeled by N.S. The legend applies to all panels. **C**: the spores in the 45%, 50% and 55% bands, obtained as described in panel A, were collected, proteins extracted, resolved by SDS-PAGE (top panel; Coomassie-stained gel) and subject to immunoblot analysis with anti-CdeM and anti-CdeC antibodies (middle and bottom panels). Black arrows show the position of CdeC or CdeM in Coomassie; red arrows show the position of multimeric species of CdeC or CdeM. The blue arrow shows the position of a species that cross-reacts with the anti-CdeM antibody. The asterisk shows the position of possible CdeM or CdeC degradation products. The position of molecular weight markers (in kDa) is shown on the left side of the panels.

### CdeM is enriched in appendage-bearing spores

We next extracted the surface proteins from WT and *cdeM* spores collected from the 45%, 50% and 55% fractions, resolved the proteins by SDS-PAGE and analyzed Coomassie stained gels, which were also subject to immunoblotting with anti-CdeM and anti-CdeC antibodies. Coomassie staining revealed a similar profile for the corresponding density fractions derived from the WT or *cdeM* samples with the exception of CdeM, absent from the mutant (Fig. 6C, top panel). Importantly, only the monomeric form of CdeM was found in the 45% fraction of WT spores; this form of the protein was clearly enriched in the 50% and 55% fractions in which multimeric forms were also detected by immunoblotting (Fig. 6C). As for CdeM, only the monomeric form of CdeC was found in the 45% fraction for both WT and *cdeM* spores (Fig. 6C; bottom panel). Importantly, multimeric forms of CdeC were found in the 50% and 55% fractions, which are enriched for appendage-bearing spores.

These results suggest that CdeC is preferentially associated with a more internal layer of this structure, upon which most of CdeM is assembled. Assembly of the exosporium around spores and/or formation of the polar appendage seems to involve multimerization of both CdeC and CdeM.

### CdeM conceals other exosporium proteins

Since treatment of purified spores with glass beads in the presence of DTT leads to the isolation of empty shells, which we posit is the exosporium, we next attempted to separate the shells from spores using density gradient centrifugation. WT and *cdeM* spore suspensions were shaken with glass beads in the presence of DTT and then applied to the top of 40-60% step gradients of metrizoic acid. For both the WT and *cdeM* samples, we found that a gradient band formed at a concentration of 42% of metrizoic acid, was enriched in phase dark structures, confirmed by SEM to be mostly empty shells although some spores were also seen (Fig. 7A). A fraction at 45% was also enriched in empty shells, but contained more spores, especially for the *cdeM* sample, and a 55% fraction contained mainly appendage-bearing spores (Fig. 7A). The material isolated from the 42% (referred to as shells) and 55% (referred to as spores) gradient fractions was analyzed by SDS-PAGE and Coomassie staining, and by immunoblotting. The profile of Coomassie-stained proteins was similar for shells derived from the WT or *cdeM* samples, except for the absence of CdeM from the latter (Fig. 7B, top panel, black arrow). Immunoblot analysis confirms the presence of multimeric forms of both CdeM and CdeC in the 42% fraction (Fig. 7B, middle panels). The BclA proteins are markers for the exosporium (Diaz-Gonzalez *et al.*, 2015, Pizarro-Guajardo *et al.*, 2014). Several forms of BclA1 are detected in coat/exosporium extracts with an anti-BclA antibody, including a form of 48 kDa and a complex of about 110 kDa (Calderon-Romero *et al.*, 2018, Pizarro-Guajardo *et al.*, 2014). Under our conditions, only the 48 kDa form of BclA1 was detected and only in the 42% fraction (shells) derived from *cdeM* spores, but not in the material obtained from WT spores (Fig. 7D, bottom panel). Both monomeric and multimeric forms of CdeC and CdeM were detected in the 55% fraction (appendage-bearing spores) but BclA1 was not detected in this fraction in spite of an overall higher protein content (Fig. 7B, right panels). Thus, BclA1 is assembled in *cdeM* spores, is enriched in the “shells” (the 42% gradient fraction), and its extractability increases in the absence of CdeM.

**Figure 7.**
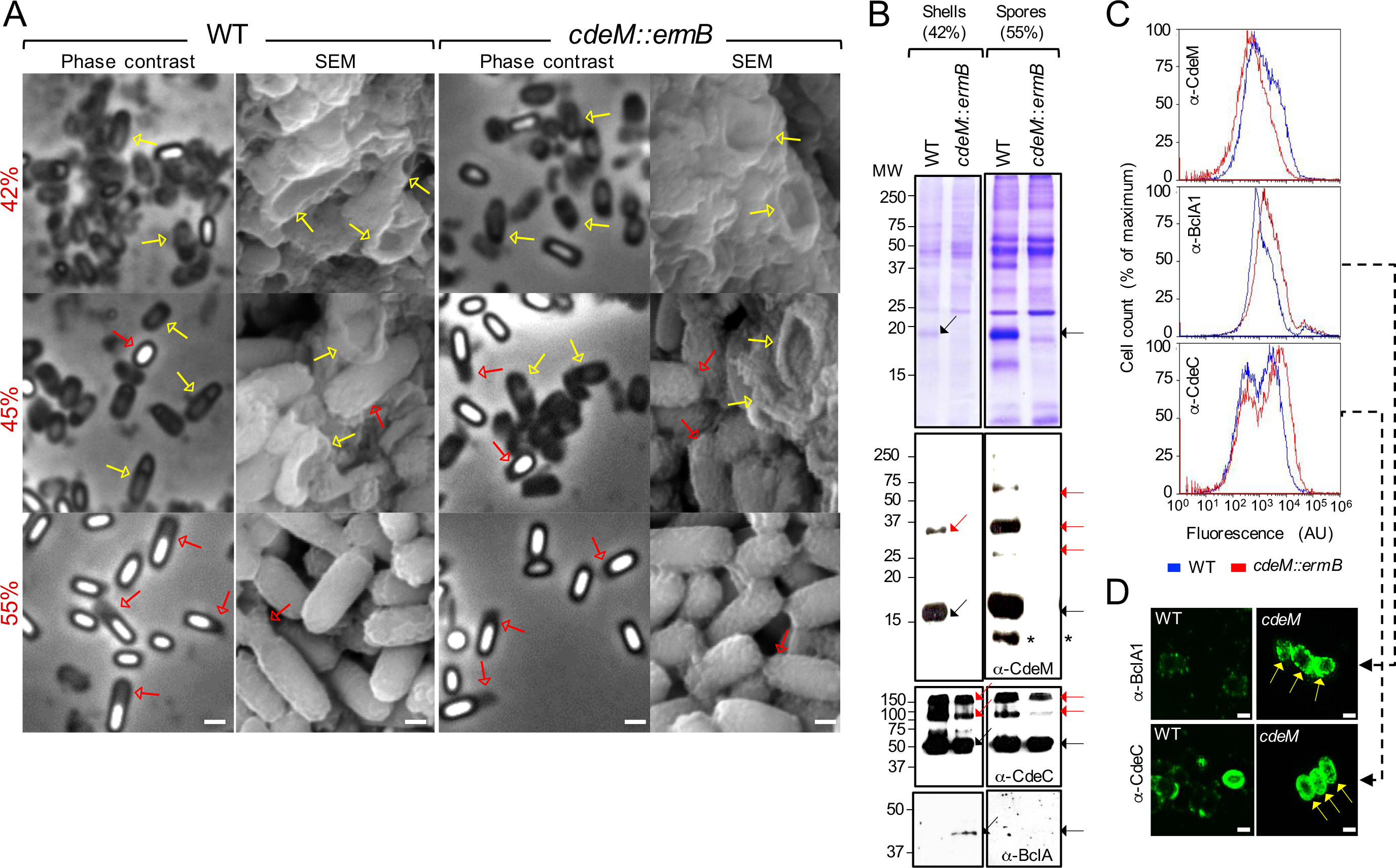
CdeM and CdeC are present in isolated exosporia. **A**: following treatment with beads in the presence of 10 mM DTT, suspensions of WT and *cdeM*::*ermB* spores were separated on step gradients of metrizoic acid. The material present in bands at 42%, 45% and 50% layers was collected and analyzed by phase contrast and SEM. Note the enrichment of empty shells in the 42% layer. The yellow arrows indicate empty exosporium shells; the red arrows indicate the position of appendage/exosporium-bearing spores, enriched in the 55% fraction. Higher magnification images of the empty shells released from WT spores are shown at the bottom. Scale bar, 1 μm. **B**: SDS-PAGE (top) and immunoblot analysis of the 42% and 55% gradient fractions with anti-CdeM, anti-CdeC and anti-BclA1 antibodies. Black arrows show the position of CdeC, CdeM or BclA1; red arrows show the position of multimeric species of CdeC or CdeM. The asterisk shows the position of a possible CdeM degradation product. Molecular weight markers (in kDa) is shown on the left side of the panels. **C**: purified WT and *cdeM*::*ermB* spores were analyzed by flow cytometry with the indicated antibodies and subject to immunofluorescence microscopy (**D**) with anti-CdeC and anti-BclA1 antibodies. The yellow arrows indicate spores with a strong fluorescence signal. AU, arbitrary units. Scale bar, 1 μm.

We used flow cytometry following immunostaining to test whether BclA1 could be more exposed in spores of the *cdeM* mutant. CdeM is detected at the surface of WT spores (Fig. 7C, top panel, blue trace), consistent with the localization studies (Fig. 3), the release of the protein following the bead/DTT treatment of spores (Fig. 5) and the enrichment of the protein in appendage-bearing spores (Fig. 6). In line with the idea that CdeM conceals BclA1, the signal for BclA increased in *cdeM* spores as compared to the WT (Fig. 7C, middle panel, red trace). Moreover, the signal for CdeC also increased in *cdeM* spores relative to the WT (Fig. 7C, red trace in the bottom panel). Fluorescence microscopy of the same spores confirms the increased decoration of *cdeM* spores by the anti-BclA1 and anti-CdeC antibodies (Fig. 7D, yellow arrows). Thus, our results suggest that in the exosporium-like layer, both BclA1 and CdeC are concealed by CdeM.

### CdeM is required for proper spore germination

The abnormal assembly of the spore surface layers in the *cdeM* mutant raised the possibility that spore germination could be affected. To test this possibility, we monitored spore germination by measuring the drop in the absorbance at 580 nm of a suspension of heat-activated spores, following addition of TA (Burns *et al.*, 2010, Sorg & Sonenshein, 2008, Wilson *et al.*, 1982). In the presence of 5% TA, but not in its absence, the OD_580_ of the heat-activated WT spore suspension dropped by about 30% of the initial in 25 min (Fig. 8A, close black symbols) whereas the OD_580_ of the *cdeM*^*C*^ spore suspension dropped 35% (close blue symbols). Germination was essentially completed 30 min after TA addition and was followed by spore outgrowth. Strikingly, for *cdeM* spores, a drop of 20% in the OD_580_ of the initial suspension takes place even in the absence of TA (Fig. 8A, open red circles) but the addition of TA caused a 50% drop in the OD_580_ in 15 min (close red circles). Thus, spores of the *cdeM* mutant seem to germinate faster and more extensively that WT or *cdeM*^*C*^ spores. As a control experiment, we used spores from a *sleC* mutant that are impaired in TA-induced germination (Adams *et al.*, 2013, Francis *et al.*, 2013) and found no germination under our conditions (Fig. 8A, green dots). Outgrowth of the *cdeM* spores followed a kinetics similar to that of WT spores (Fig. 8A). We should note that no difference in TA-induced germination was seen between WT and *cdeM* spores in a recent study (Calderon-Romero *et al.*, 2018). However, while we used 5% TA, Calderón-Romero and co-authors used 10% TA; a higher concentration of TA could potentially mask differences between WT and mutant spores. In any event, since *cdeM* spores lack the electrondense outer layer of the exosporium, increased permeability of the spore surface layers to TA may explain, at least in part, the increased germination.

**Figure 8.**
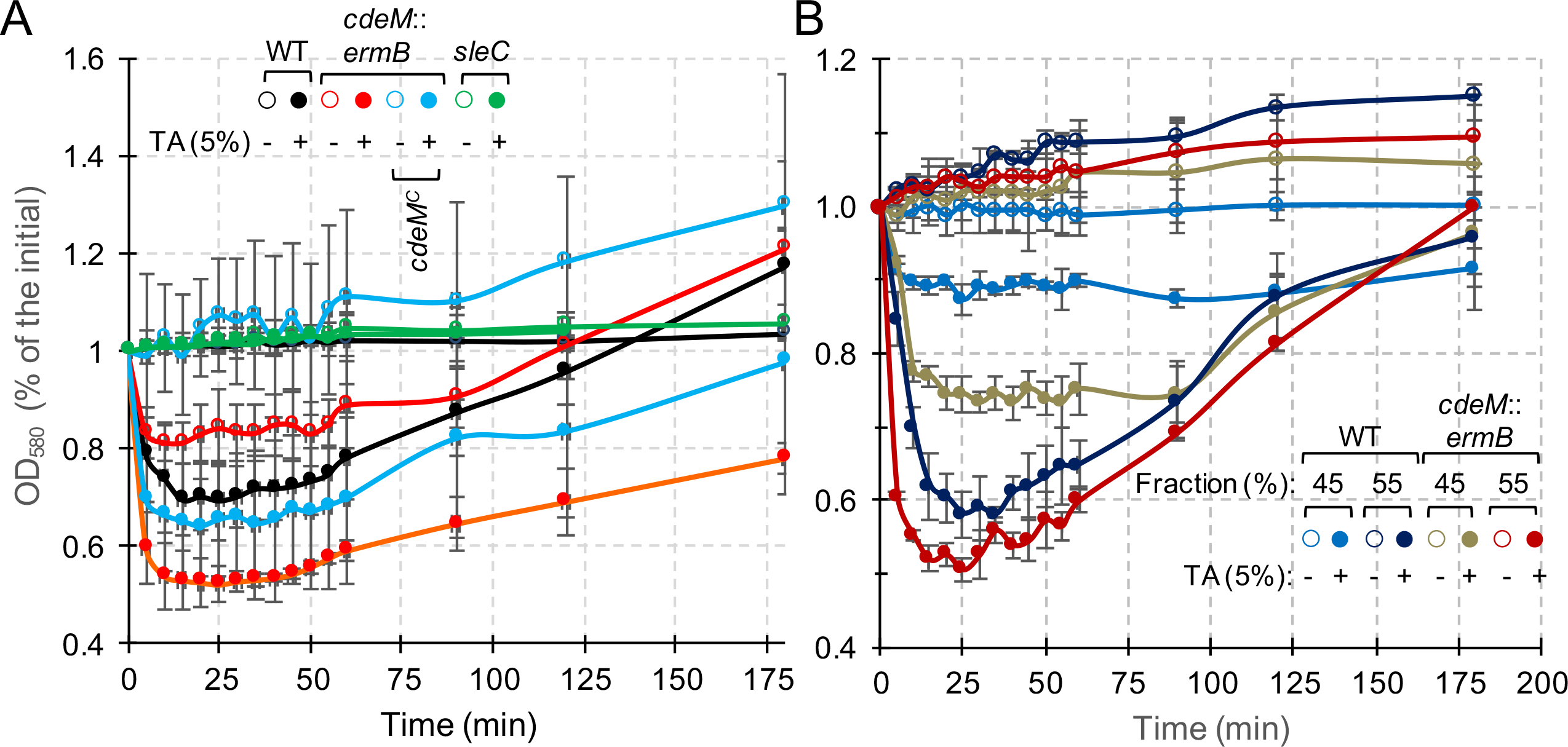
The polar appendage and CdeM have opposite effects on spore germination. **A**: Germination curves for spores of the WT strain, the *cdeM::ermB* mutant, the complementation strain *cdeM::ermB/Tn916::cdeM* and a *sleC* mutant. Purified spores were heat activated in PBS and germination induced by the addition of TA, 5% (filled symbols). In control experiments, spores were heat activated but no TA was added (open symbols). Germination was followed by the decrease in the OD_600nm_ over time at 37°C under anaerobic conditions. **B**: The kinetics of germination was monitored as described in panel A, except that the WT and *cdeM*::*ermB* spore suspensions were fractionated on gradients of metrizoic acid and the 45% (enriched in spores with no appendage) and 55% (enriched in appendage bearing spores) fractions used.

### The spore polar appendage has a direct role in germination

The *cdeM* mutant produces spores with no exosporium but that also show a short and disorganized appendage, raising the possibility that this structure contributes directly to germination. To address this question, we fractionated the population of WT and *cdeM* spores by density centrifugation and we monitored germination of the fractions enriched in smooth spores (at 45% of metrizoic acid; see also Fig. 6A and B), or in appendage-bearing spores (55% fraction). No germination was observed, for the WT or *cdeM* fractions, in the absence of TA (Fig. 8C, open symbols). In the presence of 5% TA, the smooth spores of the WT (45% fraction; Fig. 8C, close blue symbols) germinated poorly as compared to those with appendages (55% fraction; close dark blue symbols). For the *cdeM* mutant, the appendage-lacking spores (45% fraction; Fig. 8C, close green symbols) also germinated to a lower extent than the appendage-bearing spores (55% fraction; close red symbols). The appendage-bearing *cdeM* spores, however, germinated to a greater extent that the appendage-bearing WT spores and the appendage-lacking *cdeM* spores germinated better that their WT, appendage-lacking, counterparts (Fig. 8C). We infer that the appendage itself contributes directly for proper spore germination.

### *The* cdeM *mutant shows enhanced virulence*

Because spores of the *cdeM* mutant have an altered surface and germination properties, we next wanted to assess the infectivity of the mutant spores. We used the hamster model of infection in which WT spores cause a rapid and fulminant infection leading to high mortality (Goulding *et al.*, 2009, Sambol *et al.*, 2001). Five days following administration of clindamycin by gavage, a group of 6 female hamsters were challenged with WT spores, whereas 8 hamsters were challenged with *cdeM* spores, both at an infective dose of 10^3^ spores. For WT spores, the clinical end-point was reached in 7 days, with the first animals dying at day 3 of the experiment (Fig. 9A). In contrast, the clinical end-point was reached at day 5 for *cdeM* spores, with the first animals dying at day 2 (Fig. 9A). Thus, *cdeM* spores are more infective than WT spores. For both WT and *cdeM* spores, no animals survived (Fig. 9A). We were not able to purify sufficient spores of the two main appendage morphotypes for either the WT or *cdeM* mutant, and thus we were not able to directly assess the role of the appendage in virulence. Since overall, unfractionated spores of the *cdeM* mutant germinate more efficiently than WT spores in response to TA (Fig. 8A), it seems likely that the increased infectivity of *cdeM* spores is related to their increased germination (Fig. 8A; see also above).

Spores of the mutant, however, also show an altered surface which is likely to influence binding to host tissues and cells (Calderon-Romero *et al.*, 2018, Mora-Uribe *et al.*, 2016, Paredes-Sabja & Sarker, 2012) and previous work has shown that *cdeM* spores are impaired in colonization of axenic mice (Janoir *et al.*, 2014). In an attempt to determine whether the *cdeM* mutation affected the ability of spores to bind to host cells, we measured adhesion of spores to hamster caeca *ex vivo.* Isolated and washed caeca were incubated with 3x10^6^ WT or *cdeM* spores for 1, 2, or 3 h, and after washing, bound spores were enumerated by plating. Most of the WT spores bound to the hamster caecum after an incubation of 1 hour and binding did not change over time (Fig. 9B). In contrast, after 1 hour, binding of *cdeM* spores was 10-fold lower (Fig. 9B). Moreover, binding of *cdeM* spores increased over time but even after 3 hours of incubation it remained lower than for the WT (Fig. 9B). Thus, binding of *cdeM* spores to hamster caeca is impaired, in comparison to the WT spores. In a mouse colonic loop model, binding of *cdeM* spores was also impaired (Calderon-Romero *et al.*, 2018).

**Figure 9.**
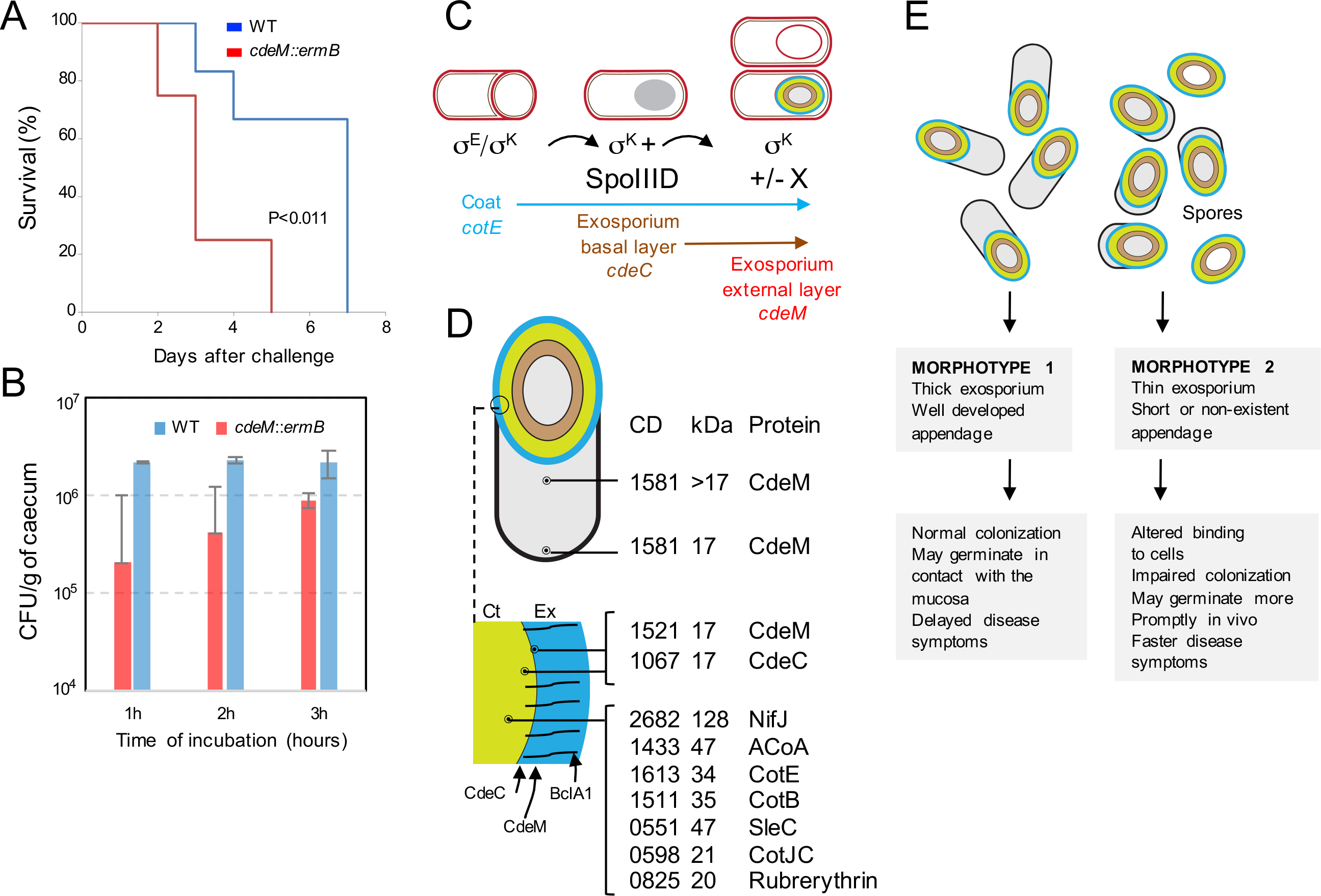
Increased virulence caused by spores of a *cdeM*::*ermB* mutant. **A**: Kaplan-Meier survival curve representing the cumulative results from two independent experiments in which clindamycin-treated Syrian golden hamsters were inoculated with WT or *cdeM*::*ermB* spores. Mean times to morbidity were: WT, 3.5 hours; *cdeM*::*ermB*, 2.6 hours (P < 0.01, log rank test). **B**: *ex vivo* adhesion of WT and *cdeM*::*ermB* spores to hamster caecum. Caeca were collected from hamsters, washed extensively, weighted, and incubated with 10^6^ WT or *cdeM*::*ermB* spores for the indicated times. After washing and grinding, binding spores were determined by plating onto blood agar plates containing taurocholic acid. The results are expressed as the number of spores per gram of ceaca. **C**: model for the regulation of assembly of the coat and exosporium of *C. difficile* spores. The onset of expression of *cotE* (prior to engulfment completion), *cdeC* (following engulfment completion when the forespore turns phase dark, under the joint control of σ^K^ and SpoIIID) and *cdeM* (when the forespore becomes phase bright) is represented as a function of the stages in morphogenesis. “X” is a putative regulatory protein, either an activator or a repressor, thought to delay expression of the σ^K^-controlled *cdeM* gene (see text for details). **D**: model for the arrangement of CdeM and CdeC in *C. difficile* spores. CdeC is represented as being part of basal layer upon which CdeM assembles conferring electrondensity to the outer layers of spores. CdeM, however, may also be located in more internal layers of the coat. In the absence of CdeM, several other proteins, including BclA1 and a CdeC, become more extractable. CdeM is found around the entire spore, but multimeric forms are enriched in the spore polar appendage. **E**: characteristics of the two main spore morphotypes (1, with a well developed appendage; 2, with a short or non-existent appendage). Spores of morphotype 1 may germinate in contact with the colonic mucosa; spores of morphotype 2 do not colonize efficiently, but may germinate more promptly in vivo and show enhanced virulence.

## Discussion

### *Expression of* cdeM *in relation to spore morphogenesis*

Our data suggests that genes coding for components of the coat and exosporium of *C. difficile* spores are expressed according to a temporal cascade that may reflect the order of assembly of the various structures. It has been shown that the genes coding for morphogenetic proteins SpoIVA and SipL are under the control of σ^E^, as are other genes coding for components of the coat layers (Fimlaid *et al.*, 2013, Pereira *et al.*, 2013, Pishdadian *et al.*, 2015, Putnam *et al.*, 2013, Saujet *et al.*, 2013). These genes are expressed mainly prior to engulfment completion, which is the main period of activity of σ^E^ and some additionally require SpoIIID (Pishdadian *et al.*, 2015). Some activity of σ^K^ is detected at this stage, but it increases following engulfment completion, explaining the early expression of *cotE* (Pereira *et al.*, 2013, Serrano *et al.*, 2016). Expression of *cdeC*, which is under the joint control of σ^K^ and SpoIIID, is only detected following engulfment completion (Pishdadian *et al.*, 2015). The onset of *cdeM* expression, which we now show to be under the direct control of σ^K^ (Fig. 2A and B) is detected later in morphogenesis, when the spore becomes phase bright. It is presently unknown how expression of *cdeM* is delayed relative to that of *cdeC.* The joint requirement for σ^K^ and SpoIIID for the expression of *cdeC* (Pishdadian *et al.*, 2015) suggests that the accumulation of SpoIIID along time may activate the expression of genes with SpoIIID-binding regions of progressively lower affinity. It is not presently known, however, whether the late expression of *cdeM* requires direct binding of SpoIIID to the promoter region of the gene. SpoIIID may only function as an activator in *C. difficile* ((Pishdadian *et al.*, 2015); but see also (Saujet *et al.*, 2013)) but in *B. subtilis* it also functions as a repressor (Eichenberger *et al.*, 2004, Halberg & Kroos, 1994, Zhang & Kroos, 1997). Thus, an alternative scenario is that the action of a *cdeM* repressor is relieved at a late stage in morphogenesis (Fig. 9C, “+/− X”, where X may be SpoIIID).

Expression of *cdeM* is only detected at high levels in 43% of the sporangia of phase bright spores, while high level expression of *cdeC* is detected in about 4% of the phase grey spores and in 24% of the sporangia of phase bright spores (Fig. 2E). Given the role of the two proteins in the assembly of the exosporium ((Calderon-Romero *et al.*, 2018); this work), it seems plausible that the two main morphotypes described herein and detected in a WT population of spores arise, at least in part, from sporangia in which the combined expression of *cdeC* and *cdeM* occurs above or below a certain threshold or does not take place. The fraction of appendage-bearing spores produced by the epidemic strain R20191 is higher than for strain 630Δ*erm* (Fig. S5) suggesting that the ratio between spores with or without an appendage is an important trait differentiating epidemic strains. The mechanism by which expression of *cdeM* is restricted to a fraction of the sporangia is unknown; it may be related to the activity of SpoIIID, as suggested above, or an as yet unknown factor (“X” in Fig. 9C). Previous work has shown that *cdeM* was the most highly expressed gene following infection of axenic mice with *C. difficile* cells (Janoir *et al.*, 2014). While suggesting that the contribution of CdeM for the assembly of the exosporium is important in the host, consistent with the colonization defect of a *cdeM* mutant (Janoir *et al.*, 2014), the factors that contribute to the differential expression of *cdeM* at least in vivo are presently unknown. Although it is tempting to conclude that most if not all of the spores that are produced in vivo have a robust appendage, it is also possible that the two sub-populations/morphotypes are maintained in vivo, because they serve different functions (see also below).

### Function of CdeM in the assembly of the exosporium

CdeM is a critical determinant of assembly of both the exosporium and a spore polar appendage ((Calderon-Romero *et al.*, 2018); this work). This suggests that CdeM is both a component of the exosporium that surrounds the spore and of the appendage (Fig. 9D), consistent with the abundance of CdeM and the localization of the protein around the spore and in the appendage as assessed both with a CdeM-SNAP fusion and immunofluorescence. A difference between the two approaches is that CdeM-SNAP labels the bulk of the appendage, whereas by immunofluorescence it is only the edge of the appendage that is labeled (Fig. 2). Possibly, CdeM is either modified in the appendage region and/or part of a tightly packed structure to which the antibodies are not accessible. Several lines of evidence suggest that CdeM undergoes disulfide cross-linking and this in turn may contribute to the rigidity of the spore surface appendage included. Firstly, CdeM is a Cys-rich and the partially purified protein forms multimers (Fig. S3); secondly, multimeric forms of CdeM are also detected in coat/exosporium extracts prepared from WT spores (Fig. 3). Finally, enrichment for an appendage-bearing sub-population of spores as well as enrichment for “shells” derived from those spores through mechanical disruption of spores in the presence of reducing agents, also enriches for multimeric forms of CdeM (and also of CdeC) (Figs. 5-7). Cross-linking of CdeM, may also confer rigidity to the spore, as indicated by our inability to produce a surface topology image of *cdeM* spores using AFM (Fig. 5).

Some of the CdeM protein is released upon treatment of the spores with beads/DTT, suggesting the existence of a sub-population more loosely attached to the exosporium and/or associated with the more internal coat layers. In contrast, no CdeC is released during isolation of the exosporium, suggesting a higher degree of cross-linking of this protein or a more internal localization within the exosporium (Fig. 5). Other lines of evidence support a more internal localization of CdeC within the exosporium. Firstly, in our hands, the assembly of CdeC is not affected by disruption of *cdeM* (Fig. 3A); a decrease in both low (44 kDa) and high (120 kDa) forms of CdeC in spores of a *cdeM* mutant, recently reported, may be due to differences in extraction conditions (Calderon-Romero *et al.*, 2018). We note for example, that under our experimental conditions, only a form of CdeC of about 50 kDa is detected in whole coat/exosporium extracts (Fig. 3), but that in other forms of CdeC are detected in enriched exosporia/shells (Fig. 7B; see bands at ≈75 kDa and ≥150 kDa). Nevertheless, a reduction in these two forms of CdeC in cdeM spores relative to the WT is in agreement with the general conclusion that CdeC and CdeM somehow influence each other (Calderon-Romero *et al.*, 2018). Secondly, CdeC becomes more accessible to antibodies in the absence of CdeM and likewise, BclA1 becomes more extractable and more accessible to antibodies in the absence of CdeM (Fig. 7B-D). Finally, intact exosporia/shells are still isolated from a *cdeM* mutant, although they are thinner (Fig. S5), consistent with a model in which CdeM assembles on top of a scaffold or basal layer of which CdeC may be part of (Fig. 9D). This putative basal layer may contain additional Cys-rich proteins such as CdeA and CdeB which have also been identified as part of the exosporium and BclA may be attached to this basal layer (Diaz-Gonzalez *et al.*, 2015, Phetcharaburanin *et al.*, 2014, Pizarro-Guajardo *et al.*, 2014). Deposition of CdeM on top of this more internal layer forms both the electrondense, surface exposed part of the exosporium and the appendage (Fig. 9D). Although the intervening proteins are not related, this model calls to mind studies in *B. cereus* and *B. anthracis.* The exosporium has a basal layer formed by several proteins, including the Cys-rich proteins CotY and CotZ, that self assemble into a crystalline hexagonal lattice through cooperative disulfide bond formation and BclA is attached to this basal layer (Ball *et al.*, 2008, Kailas *et al.*, 2011).

### Evidence for an escape route for outgrowing spores

AFM revealed the existence of a region of lower rigidity running along the long axis of WT spores. This region is most likely at the origin of the apertures seem upon treatment of intact spores with glass beads and DTT and that allow the isolation of the shells/exosporia, or at least the outermost layers of this structure. Apertures are also formed during germination of *C. sporogenes* spores, although in this case close to one of the spore pole, thought to facilitate spore emergence and cell outgrowth (Brunt *et al.*, 2015). Apertures in the exosporium are also formed during germination of spores of *B. anthracis*, which are formed at a “botllecap” structure located at one of the spore poles (Steichen *et al.*, 2007). The apertures formed in the exosporium during germination of spores of different species may form at regions of lower rigidity built into the structure during its assembly and may result, in part, from increased turgor pressure of the forespore cytoplasm (Brunt *et al.*, 2015, Steichen *et al.*, 2007). It seems likely that the apertures that result from chemical/mechanical disruption of *C. difficile* spores are also formed during spore germination.

### Biological significance of the two main spore morphotypes

Our study suggests that the two main morphotypes with respect to the assembly of the polar appendage, *i.e.*, robust appendage or disorganized or absent appendage, show different germination properties (Fig. 9E). Both WT and *cdeM* mutant spores with an appendage germinate better in response to taurocholate than those without an appendage (Fig.8). The WT spores with an appendage, however, germinate slower than their *cdeM* counterparts (Fig. 8C), possibly because in the absence of CdeM the spore is more permeable to germinants (Fig. 8A). The more permeable spores may germinate more promptly in vivo explaining why hamsters that received *cdeM* spores succumbed to infection faster than animals that received WT spores (Fig. 9A and D). Consistent with this interpretation, the overall population of *cdeM* spores germinates faster than WT spores in response to taurocholate (Fig. 8A).

The presence of the appendage may be important for germination by allowing a structural alteration of the exosporium that facilitates germination. For example, the presence of the polar appendage may facilitate opening through the region of rigidity along the spore longer axis. The appendage-bearing spores may germinate in contact with cells in vivo. Cells of a *cdeM* insertional mutant shows impaired ability to colonize axenic mice (Janoir *et al.*, 2014). The impaired colonization may result from the misassembly of the spore surface in vivo and an impaired interaction of the spores with the colonic mucosa ((Janoir *et al.*, 2014); this work). In line with this idea, spores of the *cdeM* mutant show impaired binding in a colonic loop mouse model (Calderon-Romero *et al.*, 2018), and also impaired binding to isolated hamster caeca (Fig. 9B). This is line with published work indicating that the exosporium or specific components of this structure, including CdeM, are needed for efficient spore binding to host cells ((Calderon-Romero *et al.*, 2018)and references therein).

While the expression pattern of *cdeM* may be a key factor in the production of the different spore morphotypes, as suggested above, the polymorphisms identified in *cdeM* (Fig. S4 and S7; see also the supplemental text) may also influence, at least in part, the structure of the exosporium which varies across strains (Pizarro-Guajardo *et al.*, 2016a, Pizarro-Guajardo *et al.*, 2016b). Perhaps significantly, the sequence length polymorphism at region P3 most likely eliminates the last two helices of CdeM in strain E10 (Fig. S3A and B; Fig. S7). It seems plausible that variations in the expression level and/or sequence of *cdeM*, may be related to variations in the structure of the spore surface layers and play a role in evasion of the host immune system.

## Supporting information

## Acknowledgments

We thank Ana Henriques for help with art work. This work was also financially supported by Project LISBOA-01-0145-FEDER-007660 (“Microbiologia Molecular, Estrutural e Celular”) funded by FEDER funds through COMPETE2020 - “Programa Operacional Competitividade e Internacionalizaçāo” (POCI), partially supported by project ONEIDA (LISBOA-01-0145-FEDER-016417) co-funded by FEEI - "Fundos Europeus Estruturais e de Investimento" from "Programa Operacional Regional Lisboa 2020" and by national funds from FCT - “Fundação para a Ciência e a Tecnologia”, and through FCT programme IF (IF/00268/2013/CP1173/CT0006) to M.S. FCP was the recipient of a doctoral fellowship from the FCT (SFRH/BD/45459/08).

## Material and Methods

### Strains and growth of *C. difficile*

Bacterial strains and their relevant properties are listed in Table S1. The *Escherichia coli* strain DH5α (Bethesda Research laboratories) was used for molecular cloning and BL21(DE3) (Novagen) was used for the over-production of a CdeM-His_6_ fusion (see below). Luria-Bertani medium was routinely used for growth and maintenance of *E. coli.* When indicated, ampicillin (100 μg /ml), chloramphenicol (15 μg /ml) or kanamycin (30 μg/ml) was added to the culture medium. The *C. difficile* strains used in this study are congenic derivatives of the standard laboratory strain 630Δ*erm* ((Hussain *et al.*, 2005); a gift from Nigel Minton); or strain R20291 ((McEllistrem *et al.*, 2005, Stabler *et al.*, 2009); was a gift from Daniel Paredes-Sabja). *C. difficile* strains were routinely grown anaerobically (5% H2, 15% CO2, 80% N2) at 37°C in brain heart infusion (BHI) medium (Difco) or in sporulation medium (SM), as described before (Pereira *et al.*, 2013, Saujet *et al.*, 2013). When necessary, thiamphenicol (15 μg /ml) or erythromycin (5 μg /ml) was added to *C. difficile* cultures. All plasmid constructions were carried out using standard procedures and are described in detail in the supplemental material. A list of plasmids is also included in Table S1.

### General methods

The methods used for the construction of SNAP transcriptional and translation fusions, SNAP^Cd^ labelling, phase contrast, fluorescence microscopy and image analysis, RNA extraction, quantitative RT-PCR analysis and 5′-RACE, spore trypsin digestions, KCl washes and hydrophobicity assays and for the overproduction and purification of CdeM-His_6_ for polyclonal antibody production, are given in detail in the Supplemental Material.

### Complementation analysis

To complement the *cdeM::ermB* mutation in single copy, a 1117 bp region comprising the entire coding sequence of *cdeM* (483 bp) and its expected promoter region (534 bp upstream of the *cdeM* gene) was amplified by PCR using primers PCdeM Fw/CdeM Rev (the sequence of all primers is given in Table S2). The generated PCR fragment was then cloned into the SalI and XhoI sites of pMTL84121, yielding pFT43 (Table S1). The same fragment was excised from pFT43 using SalI/HindIII restriction sites and cloned into the same sites of pSMB47, yielding pFT71. AHCD159 was created by the integration of pFT71 into the chromosomal Tn*916* locus of *B. subtilis* strain BS49. AHCD715 is a transconjugant from the mating of *B. subtilis* strain AHCD159 and *C. difficile CdeM* mutant strain, resulting in the integration of the plasmid::Tn*916* fusion of the donor strain into the *C. difficile* chromosome (see Figure S1).

### Spore production, purification and fractionation

For spore production, 5 ml of BHI media was inoculated with an isolated colony of *C. difficile* and cultured overnight at 37°C under anaerobic conditions. The next day, the overnight culture was diluted 1:100 into 100 ml of fresh BHI media and the resulting culture incubated at 37°C under anaerobic conditions for 10 days. Cells were collected by centrifugation (at 4800 × *g*, for 20 min, 4°C) resuspended in cold water and stored overnight at 4°C. The cells were then collected by centrifugation (at 4800 × *g*, for 20 min, 4°C) and the sediment resuspended in 1 ml of a 20% solution of Gastrografin (Schering). Spores were purified by applying the suspension onto a 50% layer of Gastrografin and centrifugation at 10.000 × g, for 20 min at room temperature (Henriques *et al.*, 1995). The pellet was resuspended in cold distilled water, washed 10 times with distilled water, and stored at 4°C for further use. For spore fractionation, 1 ml of a spore suspension was layered on top of step gradients of Gastrografin (from 40% to 60% in steps of 5%) and centrifuged at 10000 × g for 30 min at room temperature. Fractions (0.2 ml) were collected from the bottom and processed for scanning electron microscopy, SDS-PAGE and immunoblot analysis (below).

### Extraction, fractionation and analysis of the spore exosporium

Purified spores (10 mg dry weight) were resuspended in 500 μl of distilled water in a Pathogen Lysis Tube S (from Qiagen) and DTT added to 10 mM, final concentration. The suspension was shaken in a vortex mixer for 10 min at 10.000 rpm, at room temperature. The supernatant was then collected into a new tube and centrifuged for 10 min at 5.000 × g, at room temperature. The supernatant contains the exosporium. The sediment was washed 3 times in PBS contaiing Triton X-100 (at 0.1%). Finally, the sediment was resuspended in 50 μl of extraction buffer (0.125 mM Tris-HCl pH 6.8, 5% beta-mercaptoetanol, 2% SDS, 0.025% Bromophenol Blue, 0.5 mM DTT 5% glycerol) and boiled for 10 min. Proteins were then resolved on 15% SDS-PAGE gels and subject to immunoblot analysis with anti-CdeC, anti-CdeM or anti-BclA1 antibodies as described below.

### Spore coat extraction and mass spectrometry

Spores were resuspended in 50 μl of extraction buffer (0.125 mM Tris-HCl pH 6.8, 5% beta-mercaptoetanol, 2% SDS, 0.025% Bromophenol Blue, 0.5 mM DTT 5% glycerol) to a final OD_600nm_ of 4, and boiled as described previously (Costa *et al.*, 2006). Proteins present in the supernatant fraction were resolved on 15% SDS-PAGE gels, and visualized by Coomassie brilliant blue R-250 staining. For protein identification, protein bands were excised and digested with trypsin before analysis by matrix-assisted laser desorption ionization (MALDI) (http://www.itqb.unl.pt/Services/Analytical_Services/Mass_Spectrometry).

### Spore Germination

Density-gradient-purified spores were resuspended in PBS to a final OD_600nm_ of 1 and heat activated for 10 min at 71°C. Taurocholic acid (TA) (Sigma-Aldrich) in PBS was then added to a final concentration of 5% in TA, to induce spore germination. Germination was followed by measuring the decrease in the OD_600nm_ of the spore suspension, in a plate reader, at 37°C with agitation, until no significant changes in OD_600nm_ were detected. Spore germination was followed both in the presence of oxygen or under anaerobic conditions (5% H2, 15% CO2, 80% N2), in a glove box.

### Whole cell lysates and immunoblot analysis

Samples (10 ml) of *C. difficile* cultures were collected, and the cells collected by centrifugation (at 4800 × *g*, for 20 min, 4°C). The cell sediment was washed with phosphate-buffered saline (PBS), resuspended in 1 ml of French press buffer (10 mM Tris pH 8.0, 10 mM MgCl_2_, 0.5 mM EDTA, 0.2 mM NaCl, 10% Glycerol, 1 mM PMSF) and lysed using a French pressure cell (18000 lb/in^2^). Proteins in the total cell extracts (10 μg) or the spore coat or shell extracts prepared as described in the preceding sections were resolved on 15% SDS-PAGE gels and subject to immunoblot analysis with the following primary antibodies at the indicated dilutions: anti-CdeM, 1:10000, anti-CdeC (Fimlaid *et al.*, 2013) and anti-BclA1 (Phetcharaburanin *et al.*, 2014), both at 1:1000. A rabbit secondary antibody conjugated to horseradish peroxidase (Sigma) was used at a dilution of 1:5000. The immunoblots were developed with enhanced chemiluminescence reagents (Amersham Pharmacia Biotech).

### Immunofluorescence microscopy of sporangia and spores

Samples (500 μl) from sporulating cells were fixed with HistoChoice (from Amresco) for 15 min at room temperature. The cells were collected by centrifugation (6.000 × g, for 2 min at room temperature) the sediment resuspended in 100 μl of GTE (50 mM glucose, Tris-HCl pH 7.5, 10 mM EDTA), lysozyme added (to 2.5 mg/ml) and the suspension incubated on ice for 30 min. Following incubation, the cells were washed three times with PBS supplemented with 2% bovine serum albumin (Sigma). The primary antibodies were then added, either anti-CdeM (at a 1:1000 dilution) or anti-SNAP (from New England Biolabs, at a 1:500 dilution), and incubation continued for 1 hour. The cells were washed three times with PBS and then the Alexa Fluor 594 or Alexa Fluor 488 goat anti-rabbit IgG secondary antibody was added (Molecular probes, Invitrogen) (1:500). For immunofluorescence of spores, 100 μl of a spore suspension at an OD_580_ of about 1 was used as above, except that the lysozyme incubation step was omitted.

### Flow cytometry

For flow cytometric analysis, immune-stained spores (as above) were diluted in PBS and directly applied to a S3 cell sorter (Bio-Rad) operating a solid-state laser at 488 nm. For each sample, 30.000 events were analysed. Data containing the fluorescent signals were collected using a 525/30-bp filter, and the photomultiplier voltage was set between 400 and 550 V. Data were acquired and processed using the ProSort software package (Bio-Rad).

### Scanning electron microscopy and EDX analysis

Spores were prepared for scanning electron microscopy essentially as described (Braet *et al.*, 1997). Briefly, spores were fixed with a mixture of glutaraldehyde (2,5%) and formaldehyde (1%) in phosphate buffer (0.2 M at pH7,4) for 15 min at 22°C. The fixed spores were then washed twice in PBST (phosphate saline buffer with 0,1% tween 20) and collected by centrifugation (6000 × *g*, for 5 min at room temperature). Spores were then contrasted with 1% osmium tetroxide in water for 15 min at 22°C, washed twice in PBST and collected by centrifugation (6000 × g, 5 min). After contrasting, spores were dehydrated with a gradient of ethanol (from 50% to 100%, in 10% steps). Ethanol was then substituted by 100% acetone and 20 μl of the spore suspension was poured on a golden sputter glass slide lamella. A 50 μl drop of HDMS (hexamethyldisilazane) was applied over the spores and the lamella dried in a fume hood. Spores were then gold sputtered with 10 nm gold in an electron sputter (Sputter Coater 108auto, from Cressington). Samples were imaged using a Hitachi TM3000 scanning electron microscope (SEM), at 15 kV. Spores for energy-dispersive X-ray (EDX) spectroscopy were prepared as described above and analysed using the x-ray spectroscope Quantax 70 (Bruker Corporation) in the TM3000 SEM from Hitachi. The analysis was performed at 15 kV, for a working distance of 8,5 mm, a magnification of 18.000x and an acquisition time of 200 seconds. The elements dosed were calcium and sulphur.

### Transmission electron microscopy

Samples were collected from BHI cultures 72 hours after inoculation. Cells or density gradient-purified *C. difficile* spores (prepared as above) were processed for thin sectioning transmission electron microscopy as described before (Pereira *et al.*, 2013).

### Atomic Force Microscopy

A suspension of density gradient purified spores (3 μl; as described above) were applied to a microscope slide and left to dry. A Di (Veeco/Bruker) Dimension 3100 microscope with a Nanoscope 3a controller was used for observations. Measurements were performed in air in tapping^tm^ mode with Bruker OTESP (Olympus) tips (resonance frequency is tip dependent, but close to 280kHz). In tapping mode the tip is oscillated at high frequency touching the surface as the sample is scanned, reducing drag forces.

### Virulence assays

Two groups of adult female hamsters *(Mesocricetus auratus)* weight of 90-100g, were treated with clindamycin (50 mg/kg) five days before challenge with purified *C. difficile* spores (above). At day 0, the animals were challenged with spores as follows: group 1 (6 animals), challenged with 2 × 10^3^ wild-type spores per animal; group 2 (8 animals), challenged with 7 × 10^3^ *cdeM*::*ermB* spores per animal. The protocols used were approved by the Animal Central Department of the Paris-Sud university.

### *Ex vivo* spore binding assays

Six adult female hamsters (90-100g) were euthanized and six samples of the caeca (approximate size) removed and reversed. The caeca samples were washed 8 times in PBS. Each caeca sample was divided in two pieces and incubated with 10^6^ wild-type or *cdeM*::*ermB* spores for 1, 2 and hours, after which the sections were washed 4 times in PBS, weighted and 10 ml of PBS added. The suspension was grinded for 1 min in an Ultra-Turrax homogenizer (IKA). Serial dilutions were then plated onto Columbia blood agar plates containing cysteine (concentration?) and taurocholic acid (concentration?), in duplicate.

## References

Abecasis, A., M. Serrano, L. Alves, L. Quintais, J.B. Pereira-Leal & A.O. Henriques, (2013) A genomic signature and the identification of new endosporulaion genes. J Bacteriol 195: 2101–2115.

Abhyankar, W., A.H. Hossain, A. Djajasaputra, P. Permpoonpattana, A. Ter Beek, H.L. Dekker, S.M. Cutting, S. Brul, L.J. de Koning & C.G. de Koster, (2013) In pursuit of protein targets: proteomic characterization of bacterial spore outer layers. J Proteome Res 12: 4507–4521.

Adams, C.M., B.E. Eckenroth, E.E. Putnam, S. Doublie & A. Shen, (2013) Structural and functional analysis of the CspB protease required for Clostridium spore germination. PLoS Pathog 9: e1003165.

Aktories, K., C. Schwan & T. Jank, (2017) Clostridium difficile Toxin Biology. Annu Rev Microbiol 71: 281–307.

Ball, D.A., R. Taylor, S.J. Todd, C. Redmond, E. Couture-Tosi, P. Sylvestre, A. Moir & P.A. Bullough, (2008) Structure of the exosporium and sublayers of spores of the Bacillus cereus family revealed by electron crystallography. Mol Microbiol 68: 947–958.

Barra-Carrasco, J., V. Olguin-Araneda, A. Plaza-Garrido, C. Miranda-Cardenas, G. Cofre-Araneda, M. Pizarro-Guajardo, M.R. Sarker & D. Paredes-Sabja, (2013) The Clostridium difficile exosporium cysteine (CdeC)-rich protein is required for exosporium morphogenesis and coat assembly. J Bacteriol 195: 3863–3875.

Braet, F., R. De Zanger & E. Wisse, (1997) Drying cells for SEM, AFM and TEM by hexamethyldisilazane: a study on hepatic endothelial cells. J Microsc 186: 84–87.

Brunt, J., K.L. Cross & M.W. Peck, (2015) Apertures in the Clostridium sporogenes spore coat and exosporium align to facilitate emergence of the vegetative cell. Food Microbiol 51: 45–50.

Burns, D.A., J.T. Heap & N.P. Minton, (2010) Clostridium difficile spore germination: an update. Res Microbiol 161: 730–734.

Calderon-Romero, P., P. Castro-Cordova, R. Reyes-Ramirez, M. Milano-Cespedes, E. Guerrero-Araya, M. Pizarro-Guajardo, V. Olguin-Araneda, F. Gil & D. Paredes-Sabja, (2018) Clostridium difficile exosporium cysteine-rich proteins are essential for the morphogenesis of the exosporium layer, spore resistance, and affect C. difficile pathogenesis. PLoS Pathog 14: e1007199.

Chandrasekaran, R. & D.B. Lacy, (2017) The role of toxins in Clostridium difficile infection. FEMS Microbiol Rev 41: 723–750.

Costa, T., A.L. Isidro, C.P. Moran, Jr. & A.O. Henriques, (2006) Interaction between coat morphogenetic proteins SafA and SpoVID. J Bacteriol 188: 7731–7741.

Deakin, L.J., S. Clare, R.P. Fagan, L.F. Dawson, D.J. Pickard, M.R. West, B.W. Wren, N.F. Fairweather, G. Dougan & T.D. Lawley, (2012) The Clostridium difficile spo0A gene is a persistence and transmission factor. Infection and immunity 80: 2704–2711.

Diaz-Gonzalez, F., M. Milano, V. Olguin-Araneda, J. Pizarro-Cerda, P. Castro-Cordova, S.C. Tzeng, C.S. Maier, M.R. Sarker & D. Paredes-Sabja, (2015) Protein composition of the outermost exosporium-like layer of Clostridium difficile 630 spores. J Proteomics 123: 1–13.

Driks, A. & P. Eichenberger, (2016) The Spore Coat. Microbiol Spectr 4.

Dubberke, E.R. & M.A. Olsen, (2012) Burden of Clostridium difficile on the healthcare system. Clin Infect Dis 55 Suppl 2: S88–92.

Eichenberger, P., M. Fujita, S.T. Jensen, E.M. Conlon, D.Z. Rudner, S.T. Wang, C. Ferguson, K. Haga, T. Sato, J.S. Liu & R. Losick, (2004) The program of gene transcription for a single differentiating cell type during sporulation in Bacillus subtilis. PLoS Biol 2: e328.

Escobar-Cortes, K., J. Barra-Carrasco & D. Paredes-Sabja, (2013) Proteases and sonication specifically remove the exosporium layer of spores of Clostridium difficile strain 630. J Microbiol Methods 93: 25–31.

Fimlaid, K.A., J.P. Bond, K.C. Schutz, E.E. Putnam, J.M. Leung, T.D. Lawley & A. Shen, (2013) Global analysis of the sporulation pathway of Clostridium difficile. PLoS Genet 9: e1003660.

Fimlaid, K.A. & A. Shen, (2015) Diverse mechanisms regulate sporulation sigma factor activity in the Firmicutes. Current opinion in microbiology 24: 88–95.

Francis, M.B., C.A. Allen, R. Shrestha & J.A. Sorg, (2013) Bile acid recognition by the Clostridium difficile germinant receptor, CspC, is important for establishing infection. PLoS Pathog 9: e1003356.

Goulding, D., H. Thompson, J. Emerson, N.F. Fairweather, G. Dougan & G.R. Douce, (2009) Distinctive profiles of infection and pathology in hamsters infected with Clostridium difficile strains 630 and B1. Infection and immunity 77: 5478–5485.

Halberg, R. & L. Kroos, (1994) Sporulation regulatory protein SpoIIID from Bacillus subtilis activates and represses transcription by both mother-cell-specific forms of RNA polymerase. J Mol Biol 243: 425–436.

Henriques, A.O., B.W. Beall, K. Roland & C.P. Moran, Jr, (1995) Characterization of cotJ, a sigma E-controlled operon affecting the polypeptide composition of the coat of Bacillus subtilis spores. J Bacteriol 177: 3394–3406.

Henriques, A.O. & C.P. Moran, Jr., (2007) Structure, assembly, and function of the spore surface layers. Annu Rev Microbiol 61: 555–588.

Hong, H.A., K. Hitri, S. Hosseini, N. Kotowicz, D. Bryan, F. Mawas, A.J. Wilkinson, A. van Broekhoven, J. Kearsey & S.M. Cutting, (2017) Mucosal Antibodies to the C Terminus of Toxin A Prevent Colonization of Clostridium difficile. Infection and immunity 85.

Hussain, H.A., A.P. Roberts & P. Mullany, (2005) Generation of an erythromycin-sensitive derivative of Clostridium difficile strain 630 (630Deltaerm) and demonstration that the conjugative transposon Tn916DeltaE enters the genome of this strain at multiple sites. J Med Microbiol 54: 137–141.

Isidro, J., A. Santos, A. Nunes, V. Borges, C. Silva, L. Vieira, A.L. Mendes, M. Serrano, A.O. Henriques, J.P. Gomes & M. Oleastro, (2018) Imipenem Resistance in Clostridium difficile Ribotype 017, Portugal. Emerg Infect Dis 24: 741–745.

Janoir, C., C. Deneve, S. Bouttier, F. Barbut, S. Hoys, L. Caleechum, D. Chapeton-Montes, F.C. Pereira, A.O. Henriques, A. Collignon, M. Monot & B. Dupuy, (2014) Adaptive strategies and pathogenesis of Clostridium difficile from in vivo transcriptomics. Infection and immunity 81: 3757–3769.

Jiang, S., Q. Wan, D. Krajcikova, J. Tang, S.B. Tzokov, I. Barak & P.A. Bullough, (2015) Diverse supramolecular structures formed by self-assembling proteins of the Bacillus subtilis spore coat. Mol Microbiol 97: 347–359.

Jones, A.M., E.J. Kuijper & M.H. Wilcox, (2013) Clostridium difficile: a European perspective. J Infect 66: 115–128.

Joshi, L.T., D.S. Phillips, C.F. Williams, A. Alyousef & L. Baillie, (2012) Contribution of spores to the ability of Clostridium difficile to adhere to surfaces. Appl Environ Microbiol 78: 7671–7679.

Kailas, L., C. Terry, N. Abbott, R. Taylor, N. Mullin, S.B. Tzokov, S.J. Todd, B.A. Wallace, J.K. Hobbs, A. Moir & P.A. Bullough, (2011) Surface architecture of endospores of the Bacillus cereus/anthracis/thuringiensis family at the subnanometer scale. Proc Natl Acad Sci U S A 108: 16014–16019.

Kociolek, L.K. & D.N. Gerding, (2016) Breakthroughs in the treatment and prevention of Clostridium difficile infection. Nat Rev Gastroenterol Hepatol 13: 150–160.

Lawley, T.D., N.J. Croucher, L. Yu, S. Clare, M. Sebaihia, D. Goulding, D.J. Pickard, J. Parkhill, J. Choudhary & G. Dougan, (2009) Proteomic and genomic characterization of highly infectious Clostridium difficile 630 spores. J Bacteriol 191: 5377–5386.

Lawson, P.A., D.M. Citron, K.L. Tyrrell & S.M. Finegold, (2016) Reclassification of Clostridium difficile as Clostridioides difficile (Hall and O’Toole 1935) Prevot 1938. Anaerobe 40: 95–99.

Lessa, F.C., Y. Mu, W.M. Bamberg, Z.G. Beldavs, G.K. Dumyati, J.R. Dunn, M.M. Farley, S.M. Holzbauer, J.I. Meek, E.C. Phipps, L.E. Wilson, L.G. Winston, J.A. Cohen, B.M. Limbago, S.K. Fridkin, D.N. Gerding & L.C. McDonald, (2015) Burden of Clostridium difficile infection in the United States. N Engl J Med 372: 825–834.

McEllistrem, M.C., R.J. Carman, D.N. Gerding, C.W. Genheimer & L. Zheng, (2005) A hospital outbreak of Clostridium difficile disease associated with isolates carrying binary toxin genes. Clin Infect Dis 40: 265–272.

McKenney, P.T., A. Driks & P. Eichenberger, (2013) The Bacillus subtilis endospore: assembly and functions of the multilayered coat. Nat Rev Microbiol 11: 33–44.

McKenney, P.T., A. Driks, H.A. Eskandarian, P. Grabowski, J. Guberman, K.H. Wang, Z. Gitai & P. Eichenberger, (2010) A distance-weighted interaction map reveals a previously uncharacterized layer of the Bacillus subtilis spore coat. Curr Biol 20: 934–938.

Mora-Uribe, P., C. Miranda-Cardenas, P. Castro-Cordova, F. Gil, I. Calderon, J.A. Fuentes, P.I. Rodas, S. Banawas, M.R. Sarker & D. Paredes-Sabja, (2016) Characterization of the Adherence of Clostridium difficile Spores: The Integrity of the Outermost Layer Affects Adherence Properties of Spores of the Epidemic Strain R20291 to Components of the Intestinal Mucosa. Front Cell Infect Microbiol 6: 99.

Panessa-Warren, B.J., G.T. Tortora & J.B. Warren, (1997) Exosporial membrane plasticity of Clostridium sporogenes and Clostridium difficile. Tissue Cell 29: 449–461.

Paredes-Sabja, D. & M.R. Sarker, (2012) Adherence of Clostridium difficile spores to Caco-2 cells in culture. J Med Microbiol 61: 1208–1218.

Pereira, F.C., L. Saujet, A.R. Tome, M. Serrano, M. Monot, E. Couture-Tosi, I. Martin-Verstraete, B. Dupuy & A.O. Henriques, (2013) The spore differentiation pathway in the enteric pathogen Clostridium difficile. PLoS Genet 9: e1003782.

Permpoonpattana, P., J. Phetcharaburanin, A. Mikelsone, M. Dembek, S. Tan, M.C. Brisson, R. La Ragione, A.R. Brisson, N. Fairweather, H.A. Hong & S.M. Cutting, (2013) Functional characterization of Clostridium difficile spore coat proteins. J Bacteriol 195: 1492–1503.

Permpoonpattana, P., E.H. Tolls, R. Nadem, S. Tan, A. Brisson & S.M. Cutting, (2011) Surface layers of Clostridium difficile endospores. J Bacteriol 193: 6461–6470.

Phetcharaburanin, J., H.A. Hong, C. Colenutt, I. Bianconi, L. Sempere, P. Permpoonpattana, K. Smith, M. Dembek, S. Tan, M.C. Brisson, A.R. Brisson, N.F. Fairweather & S.M. Cutting, (2014) The spore-associated protein BclA1 affects the susceptibility of animals to colonization and infection by Clostridium difficile. Mol Microbiol 92: 1025–1038.

Pishdadian, K., K.A. Fimlaid & A. Shen, (2015) SpoIIID-mediated regulation of sigmaK function during Clostridium difficile sporulation. Mol Microbiol 95: 189–208.

Pizarro-Guajardo, M., P. Calderon-Romero, P. Castro-Cordova, P. Mora-Uribe & D. Paredes-Sabja, (2016a) Ultrastructural Variability of the Exosporium Layer of Clostridium difficile Spores. Appl Environ Microbiol 82: 2202–2209.

Pizarro-Guajardo, M., P. Calderon-Romero & D. Paredes-Sabja, (2016b) Ultrastructure Variability of the Exosporium Layer of Clostridium difficile Spores from Sporulating Cultures and Biofilms. Appl Environ Microbiol 82: 5892–5898.

Pizarro-Guajardo, M., V. Olguin-Araneda, J. Barra-Carrasco, C. Brito-Silva, M.R. Sarker & D. Paredes-Sabja, (2014) Characterization of the collagen-like exosporium protein, BclA1, of Clostridium difficile spores. Anaerobe 25: 18–30.

Putnam, E.E., A.M. Nock, T.D. Lawley & A. Shen, (2013) SpoIVA and SipL are Clostridium difficile spore morphogenetic proteins. J Bacteriol.

Rabi, R., L. Turnbull, C.B. Whitchurch, M. Awad & D. Lyras, (2017) Structural Characterization of Clostridium sordellii Spores of Diverse Human, Animal, and Environmental Origin and Comparison to Clostridium difficile Spores. mSphere 2.

Rupnik, M., M.H. Wilcox & D.N. Gerding, (2009) Clostridium difficile infection: new developments in epidemiology and pathogenesis. Nat Rev Microbiol 7: 526–536.

Sambol, S.P., J.K. Tang, M.M. Merrigan, S. Johnson & D.N. Gerding, (2001) Infection of hamsters with epidemiologically important strains of Clostridium difficile. J Infect Dis 183: 1760–1766.

Samsonoff, W.A., T. Hashimoto & S.F. Conti, (1971) Appendage development in Clostridium bifermentans. J Bacteriol 106: 269–275.

Saujet, L., F.C. Pereira, M. Serrano, O. Soutourina, M. Monot, P.V. Shelyakin, M.S. Gelfand, B. Dupuy, A.O. Henriques & I. Martin-Verstraete, (2013) Genome-wide analysis of cell type-specific gene transcription during spore formation in Clostridium difficile. PLoS Genet 9: e1003756.

Serrano, M., N. Kint, F.C. Pereira, L. Saujet, P. Boudry, B. Dupuy, A.O. Henriques & I. Martin-Verstraete, (2016) A Recombination Directionality Factor Controls the Cell Type-Specific Activation of sigmaK and the Fidelity of Spore Development in Clostridium difficile. PLoS Genet 12: e1006312.

Setlow, P., (2014) Spore Resistance Properties. Microbiol Spectr 2.

Smits, W.K., D. Lyras, D.B. Lacy, M.H. Wilcox & E.J. Kuijper, (2016) Clostridium difficile infection. Nat Rev Dis Primers 2: 16020.

Sorg, J.A. & A.L. Sonenshein, (2008) Bile salts and glycine as cogerminants for Clostridium difficile spores. J Bacteriol 190: 2505–2512.

Stabler, R.A., M. He, L. Dawson, M. Martin, E. Valiente, C. Corton, T.D. Lawley, M. Sebaihia, M.A. Quail, G. Rose, D.N. Gerding, M. Gibert, M.R. Popoff, J. Parkhill, G. Dougan & B.W. Wren, (2009) Comparative genome and phenotypic analysis of Clostridium difficile 027 strains provides insight into the evolution of a hypervirulent bacterium. Genome Biol 10: R102.

Steichen, C.T., J.F. Kearney & C.L. Turnbough, Jr., (2007) Non-uniform assembly of the Bacillus anthracis exosporium and a bottle cap model for spore germination and outgrowth. Mol Microbiol 64: 359–367.

Stewart, G.C., (2015) The Exosporium Layer of Bacterial Spores: a Connection to the Environment and the Infected Host. Microbiol Mol Biol Rev 79: 437–457.

Waller, L.N., N. Fox, K.F. Fox, A. Fox & R.L. Price, (2004) Ruthenium red staining for ultrastructural visualization of a glycoprotein layer surrounding the spore of Bacillus anthracis and Bacillus subtilis. J Microbiol Methods 58: 23–30.

Wilson, K.H., M.J. Kennedy & F.R. Fekety, (1982) Use of sodium taurocholate to enhance spore recovery on a medium selective for Clostridium difficile. J Clin Microbiol 15: 443–446.

Yutin, N. & M.Y. Galperin, (2013) A genomic update on clostridial phylogeny: Gram-negative spore formers and other misplaced clostridia. Environ Microbiol 15: 2631–2641.

Zhang, B. & L. Kroos, (1997) A feedback loop regulates the switch from one sigma factor to the next in the cascade controlling Bacillus subtilis mother cell gene expression. J Bacteriol 179: 6138–6144.

