## Supplementary material for "Structure and assembly of a *Clostridioides difficile* spore polar appendage"

[1]Instituto de Tecnologia Química e Biológica António Xavier, Universidade Nova de Lisboa, Oeiras, Portugal, [2]Laboratório de Bromatologia e Defesa Biológica do Exército, Centro de Investigação da Academia Militar, Avenida Doutor Alfredo Bensaúde, 1849-012, Lisboa, Portugal; [3]Laboratoire Pathogenèse des Bactéries Anaérobies, Institut Pasteur, Paris, France, [4]Université de Paris-Sud, Faculté de Pharmacie, Département de Microbiologie, Châtenay-Malabry, France, [5]Instituto de Ciência e Engenharia de Materiais e Superfícies, Instituto Superior Técnico, Portugal; Instituto Gulbenkian de Ciência, Oeiras, Portugal[6]; [7]Université Paris 7-Denis Diderot, 75205 Paris, France.

[$]Present address: Laboratoire Pathogenèse des Bactéries Anaérobies, Institut Pasteur, Paris, France.

[#]Present address: University of Vienna, Department of Microbiology and Ecosystem Science, Division of Microbial Ecology, Vienna, Austria

[&]Present address: Roche, Basel, Switzerland.

[€]Present address: Eliho Bioscience, Paris, France.

### Supplemental Material and Methods

**Complementation of *cdeM*::*ermB* in single copy using Tn*916*.** To complement the *C. difficile cdeM* mutant (Table S1) in single copy, a 1117 bp region comprising the entire coding sequence of *cdeM* (CD630_1581; 483 bp in length) and its regulatory region (534 bp upstream of the *cdeM* start codon) was amplified by PCR using primers Psp17Fw/Sp17Rev (Table S2). The resulting PCR fragment was then cloned into the SalI and XhoI sites of pMTL84121, yielding pFT43 (Table S1). The same fragment was excised from pFT43 using SalI/HindIII and cloned between the same sites of pSMB47, yielding pFT71 (Table S1 and Fig. S2). AHCD159 was created by the integration of pFT71 into the chromosomal Tn*916* locus *of B. subtilis* strain BS49 (Fig. S2). AHCD715 is a transconjugant from the mating of *B. subtilis* AHCD159 with the *C. difficile cdeM*::*ermB* mutant, resulting in the integration of the plasmid::Tn916 fusion of the donor strain into the *C. difficile* chromosome (Fig. S1).

***SNAP$^{Cd}$* fusions.** To construct *cdeM*, *cotE* and *cdeC* transcriptional fusions to the *SNAP$^{Cd}$* reporter (Pereira *et al.*, 2013, Saujet *et al.*, 2013), the promoter regions of these genes were PCR-amplified from genomic DNA of *C. difficile* 630Δ*erm* using primer pairs Psp17-SalI Fw/Psp17 Rev, PcotE-EcoRI Fw/PcotE Rev and PcdeC-EcoRI Fw/PcdeC Rev to produce 546, 310 and 327 bp products, respectively. The *SNAP$^{Cd}$* gene was PCR amplified from pFT47 using primer pairs Psp17_SNAPFw/SNAP-HindIII Rev, PcotE_SNAP Fw/SNAP-HindIII Rev and PcdeC_SNAP Fw/SNAPHindIIIRev. The two pieces of each transcriptional fusion were joined, respectively, into a 1101 bp, 877 bp and 882 bp fragments by SOE PCR and cloned between the SalI and HindIII sites (in the case of the $P_{cdeM}$- *SNAP$^{Cd}$* fusion) and between the EcoRI and HindIII sites (for $P_{cotE}$- *SNAP$^{Cd}$* and $P_{cdeC}$-*SNAP$^{Cd}$* fusions) of pMTL84121 to produce pCAF5, pCAF7 and pCAF8, respectively. The absence of unwanted mutations was verified by sequencing the insert in all the plasmids. Plasmids pCAF5, pCAF7 and pCAF8 were introduced into *E. coli* HB101 (RP4) and then transferred to *C. difficile* 630Δ*erm* and *sigK*::*ermB* (AHCD535) strains by conjugation ((Pereira *et al.*, 2013); Table S1).

**RNA extraction, quantitative RT-PCR analysis, and 5´-RACE.** Total RNA from the strain 630Δ*erm* and the *sigF*, *sigE, sigG* and *sigK* mutants was extracted as described before (Saujet *et al.*, 2013). Briefly, cells grown in SM were harvested after 14 h (630Δ*erm*, *sigF* and *sigE*), 19 h (630Δ*erm*, *sigG*) or 24 h (630Δ*erm* , *sigK*) of growth. After centrifugation, the culture pellets were resuspended in RNApro™ solution (MP Biomedicals) and RNA extracted using the FastRNA Pro Blue Kit, according to the manufacturer's instructions. For quantitative RT-PCR experiments, 1 µg of total RNA was heated at 70°C for 10 min along with 1 µg of hexamer oligonucleotide primers $p(dN)_6$ (Roche). After slow cooling, cDNAs were synthesized for 2 h at 37°C with AMV Reverse Transcriptase (Promega), 20 mM dNTP mix and 40 U RNasin (Promega). The reverse transcriptase was inactivated by incubation at 85°C for 5 min. Real-time quantitative RT-PCR was performed twice in a 20 µl reaction volume containing 20 ng of cDNAs, 10 µl of FastStart SYBR Green Master mix (ROX, Roche) and 200 nM gene-specific primers (Table S2). Amplification and detection were performed as previously described (Saujet *et al.*, 2011). In each sample, the quantity of cDNAs of a gene was normalized to the quantity of cDNAs of the DNApolIII gene. The relative transcript changes were calculated using the $2^{-\Delta\Delta Ct}$ method (Saujet *et al.*, 2011).

5' RACE experiments were done using the system for Rapid Amplification of cDNA Ends kit (Invitrogen) according to the manufacturer's instructions. Briefly, a first strand cDNA was synthesised from total RNA extracted after 10 h of growth in TY using Superscript II and the primer LS138 (cccagtatttttctgcttctt) annealing in *CD1581*. A C-tail was then added to the 3' end of the cDNA using TdT and dCTP. The resulting fragment was amplified by the Abridged Anchor Primer and LS139 (tttctcttatctcagcttctct) annealing in *CD1581*. The resulting fragment was then sequenced by LS140 (tagctgtagattctcctctttc) annealing in *CD1581* in order to define the transcriptional start site mapping.

**Trypsin digestion of purified spores.** Purified spores (1 $OD_{580}$ unit) were incubated for 1h at 37ºC with or without trypsin (100µg/ml; Animed). The reaction was stopped by addition of SDS loading dye and the mixture boiled. The extracted coat proteins were resolved by SDS-PAGE (15%).

**Spore washes with KCl**. The method was described by Serrano et al. (1999), and was used with minor modifications. Briefly, purified spores (1 $OD_{580}$ unit) in a volume of 20 µl were incubated with 1 ml of KCl for 1 hour with orbital shaking (1 hour at room temperature). The spores were collected by centrifugation (3.000 x g, for 5 min at room temperature) resuspended in the initial volume of SDS-loading dye. Proteins in the supernatant were precipitated with and equal volume of TCA, washed two time with 1ml of 70% ethanol, and the sediment resuspend in 20 µl of SDS-loading dye. The resuspended spore and protein sediment were boiled for 8 min and the mixture resolved by SDS-PAGE (15%).

**CdeM-$His_6$ overproduction purification and polyclonal antibody production.** A DNA fragment encoding the *cdeM* gene was generated by PCR from *C. difficile* 630Δ*erm* genomic DNA using primers CdeM-pET33b Fw/ CdeM-pET33b Rev. The resulting DNA fragment was cut with NcoI and XhoI and cloned between the same sites of pET33b (Novagen) to produce pTV1. Plasmid pTV1, which codes for a CdeM-$His_6$ fusion protein, was introduced into BL21 (DE3) cells. For inducing production of

CdeM-$His_6$, IPTG (to 1 mM) was added to LB cultures at an $OD_{600nm}$ of about 0.5 and incubation continued for 4h. The cells were then harvested by centrifugation (4000 x *g*, for 10 min, at 4ºC) and the sediment resuspended in lysis buffer (20mM phosphate pH7.4, 1 mM PMSF, 10 mM Imidazole). The suspension was lysed using a French pressure cell (at 18000 $lb/in^2$) and the lysate cleared by centrifugation (15000 x *g*, 30 min at at 4ºC). Since CdeM-$His_6$ was largely insoluble, the sediment was solubilized with 8M Urea for 30 min, and the supernantant from the solubilized fraction was loaded onto a 1 ml Histrap column (Amersham Phamarcia Biotech). The bound protein was eluted with a discontinuous imidazole gradient and the fractions containing CdeM-$His_6$ were identified by SDS-PAGE. For the production of a rabbit polyclonal antibody, CdeM-$His_6$ was purified, about 200 µg resolved by SDS-PAGE (15% gels) and bands excised. The antibody was produced by Eurogentec (Seraing, Belgium).

**Spore hydrophobicity assays.** Hydrophobicity of *C. difficile* spores was determined using the bacterial adherence to hydrocarbons (BATH) method (Rosenberg, 1984). Briefly, density-gradient-purified spores were suspended in 1 ml sterile distilled water to a final $OD_{440nm}$ of about 0.5 and mixed with 100 µl of hexadecane (Sigma-Aldrich). Adherence to hydrocarbon was measured by quantifying the drop of $OD_{440nm}$ of the aqueous solution. The percent decrease in $OD_{440nm}$ for the aqueous solution was calculated according to the formula: $100 \times (OD_0-OD_f)/OD_0$, where $OD_0$ and $OD_f$ refer to the initial and final optical densities of the aqueous phase.

**$SNAP^{Cd}$ labelling, phase contrast, fluorescence microscopy and image analysis**. For SNAP labeling, cells in culture samples were mixed with the TMR-Star substrate (New England Biolabs) at a final concentration of 250 nM, for 30 min in the dark. Following labeling, the cells were collected by centrifugation (4000 x g for 5 min at room temperature), washed four times with 1 ml of PBS, and finally resuspended in

0.5 ml of PBS containing the membrane dye Mitotracker Green (MTG, 0.5 ng/ml) (Molecular probes, Invitrogen). For phase contrast and fluorescence microscopy, cells were mounted on 1.7% agarose coated glass slides and observed on a Leica DM6000B microscope equipped with a phase contrast Uplan F1 100x objective and captured with a CCD Andor Ixon camera (Andor Technologies). Images were acquired and analyzed using the Metamorph software suite (version 5.8; Universal Imaging) and adjusted and cropped using *ImageJ* (http://rsbweb.nih.gov/ij/).

**Protein taxonomic distribution and polymorphism analysis.** Protein homology search was performed using *phmmer* implemented in the HMMER web server (Finn *et al.*, 2015). This algorithm is analogous to BLASTP as it takes a single protein sequence and queries it against a target sequence database for similar sequences. HMMER, however, uses a probabilistic model especially efficient in the detection of distant homologues (Finn *et al.*, 2011). We query the CdeM sequence from *C. difficile* str. strain 630 against both UniProtKB (version 2017_02, downloaded on 2017-02-03; http://www.uniprot.org/) and *Ensemble Bacteria* (version 34, downloaded on 2017-02-03; http://bacteria.ensembl.org/index.html) sequence databases, and used an *E*-value threshold of $10^{-4}$ and default search parameters. We extracted all significant hits and aligned them in MAFFT v. 7.154, using the G-INS-I method and default parameter values (Katoh & Standley, 2013). Phylogenetic analyses were carried out with RaxML v. 8.0.26 (Stamatakis, 2014) with HIVb+G model of evolution that was determined with Prottest version 3.4 (Darriba *et al.*, 2011). Nodal support was estimated with nonparametric bootstrap analysis using an automatic frequency-based criterion (autoFC option) to determine the number of replicates.

The alignment of selected protein sequences were visualized using JalView (Waterhouse *et al.*, 2009). Several statistics were estimated using available tools on

Jalview (Waterhouse *et al.*, 2009). Alignment conservation measures the number of conserved physico-chemical properties for each column of the alignment. This calculation is based on the AMAS method (Livingstone & Barton, 1993). Alignment quality is an ad-hoc measure of the likelihood of observing the mutations (if any) in a particular column of the alignment. The consensus is the percentage of the modal residue per column. The last nine rows below the alignment are the results from the JNet secondary structure prediction tool that uses the Jpred Server version 4.0.0 (Cuff & Barton, 1999, Drozdetskiy *et al.*, 2015). We used the full sequence alignment for a JNet prediction on the sequence *C. difficile* str. 630. The annotation bars are as follows (Fig. S3): Lupas_21, Lupas_14, and Lupas_28 are coiled-coil predictions for three windows sizes (window size of 14, 21 and 28); JNetPRED is the consensus prediction were red tubes indicates $\alpha$-helices. JNetCONF is the confidence estimate for the prediction. JNetHMM is the HMM profile based prediction; JNETPSSM is the PSSM based prediction; JNETJURY indicates with an '*' whether the JNETJURY was invoked to rationalize significantly different primary predictions; and JNet burial is the JNet burial prediction.

### Supplemental Results and Discussion

#### The *cdeM*::*ermB* mutation affects the surface properties of spores

The impaired assembly of the surface layers of the spore in the *cdeM* mutant is reminiscent of the phenotype of *cotE* mutants of *B. subtilis*; spores produced by a *cotE* mutant lack the outer coat and crust layers (McKenney *et al.*, 2010, Zheng *et al.*, 1988) and are sensitive to cortex lytic enzymes such as lysozyme (Costa *et al.*, 2007, Zheng *et al.*, 1988). We reasoned that the impaired assembly of the spore surface layers could likewise render spores of the *cdeM* mutant sensitive to lysozyme. However, and in agreement with previous work (Calderon-Romero *et al.*, 2018)) we did not detect any significant change in the ability of *cdeM::ermB* spores to resist to lysozyme, as compared to WT spores. Also, no differences in heat resistance were observed for *cdeM::ermB* spores when compared to the WT (Calderon-Romero *et al.*, 2018).

Hydrophobic interactions have been shown to be important for different aspects of pathogenesis, either directly, like the adherence to host epithelial cells (Andersson & Freeman, 1998, Bozue *et al.*, 2007, Paredes-Sabja & Sarker, 2012), or indirectly, like the adherence to surfaces, making decontamination difficult (Joshi *et al.*, 2012). In the pathogen *Bacillus anthracis*, BclA, a major component of the spore exosporium, affects spore hydrophobicity and contributes to increased adherence of the spore to other types of cells beyond macrophages, (Bozue *et al.*, 2007, Brahmbhatt *et al.*, 2007). In *C. difficile* it has also been shown that the spore surface layers, such as the exosporium layer, significantly contribute to the *C. difficile* spore hydrophobicity (Escobar-Cortes *et al.*, 2013). Since CdeM is a determinant for the assembly of the spore outer coat, we evaluated the impact of *cdeM* disruption on spore hydrophobicity, accessed using a BATH assay (Paredes-Sabja & Sarker, 2012). Results showed that 39% of *cdeM::ermB* spores were present in the hydrocarbon organic phase, contrasting with 52% of the WT spores. Complementation of the *cdeM* mutation with a single copy of *cdeM* (inserted into Tn*916*) restored the WT phenotype (50% of spores in the organic phase). The results show that CdeM is required for normal spore hydrophobicity, in line with its role in proper assembly of the spore surface layers.

**The polar appendage in spores of the epidemic strain R20291**

To determine whether the bifurcation of a WT population into spores with a well organized appendage (type A) and a disorganized appendage (type B) was also seen for an epidemic strain, we measured the percentage of the two types of spores for strain R20291, of ribotype 027 (Stabler *et al.*, 2009). We found that type A appendages represented 50% of the spores whereas type B appendages represented 10% of the population (Fig. S5A). Thus, at least for this strain, the percentage of type A appendages in the population was higher than for strain 630Δ*erm* (Fig. 4; see also the main text).

Spores produced by strain R20291 were wider than 630Δ*erm* spores (Fig. S5B; average width of 0.78±0.09 µm and 0.67±0.07 µm, respectively), consistent with the presence of a more robust exosporium in spores of the first (Pizarro-Guajardo *et al.*, 2016). They were also significantly longer than 630Δ*erm* spores, as measured in the phase contrast images (Fig. S5B; average length from Ph. contrast: 1.53±0.25 µm vs.1.30±0.21 µm; ). Differences in spore length were also detected in the FM4-64 images, but are less pronounced than the differences observed by phase contrast (Fig. S5B; average length from FM4-64: 1.11±0.15 µm vs 1.03±0.25 µm). The larger spore length as measured in the phase contrast images relative to 630Δ*erm* spores, together with the presence of a thicker exosporium in R20191 spores (Pizarro-Guajardo et al., 2016a,b), is consistent with the view that the appendage is an integral part of this structure.

We also wanted to determine whether the beads/DTT treatment of R20291 spore suspensions produced empty shells as found for spores of strain 630Δerm. The beads/DTT treatment of R20291 spores produced empty shells which showed a bumpy appearance (Fig. S5C). The exosporium of R20291 spores is thicker than that of 630Δ*erm* spores, and shows a convoluted (or bumpy) appearance in TEM images (Pizarro-Guajardo *et al.*, 2016, Rabi *et al.*, 2017). The observation that the empty shells derived from spores of strain R20191 had a bumpy appearance lends support to our contention that the shells represent the isolated exosporium. Importantly, the thickness of the shell, measured in field emission SEM images at the edge of the apertures produced by the beads/DTT treatment, was of 0.091±0.001 μm for 630Δ*erm* spores, and of 0.14±0.024 μm for R20191 spores (Fig. S5). This is in agreement with the presence of a thicker exosporium in spores of the latter strain.

**CdeM is polymorphic**

A search performed with the sequence of CdeM from *C. difficile* strain 630 against either UniProtKB and Ensemble Bacteria databases retrieved 229 different genomes of *C. difficile* and only of this organism. Because of the extensive taxonomic representation of genomes in those databases (4542 and 41610 bacterial genomes, respectively), together with the probabilistic model used for homology detection, we are confident that if homologues sequences were present in other sequenced genomes we would have been able to detect them. We infer, therefore, that CdeM is unique to *C. difficile*, a conclusion also presented in a recent study (Calderon-Romero *et al.*, 2018) and most likely present in all sequenced strains of this organism (Fig. S9). Figure S3 shows the alignment of a subset of those proteins that were selected to maximize intraspecific polymorphism (see also Table S3). In general, CdeM is highly conserved among strains. CdeM shows, however, four regions of sequence polymorphism (*i.e.*, sequence variation at the intraspecific level), P1 through P4. Both substitutions (region P1, P2 and P4) and sequence length polymorphism (region P3) are found (Fig. S3). The large majority of the homologous sequences present on the *Ensembl Bacteria* database are restricted to two main CdeM sequence types: type A in strain 630 (in 150 out of 229 genomes) and type C in strain E24 (represented in 45 genomes) (Fig. S9 and Table S4). These two sequence types are closely related and differ mainly on the polymorphic region closest to the N-terminus of CdeM (region P1 in Fig. S3B). Secondary structure predictions suggest that CdeM is formed by a series of $\alpha$-helices with relatively short sequences in between the helices (Fig. S3A). The N-terminal end of the protein (residues 1-20) appears to be unstructured, as well as the C-terminal end of the protein (residues 150-173), which has a Cys-rich motif (Fig. S3B; residues 150-160, **CC**HK**C**HK**C**N**C**N**CC**RK, in strain 630). While regions P1, P2 and P4 are outside the predicted helices, the sequence length polymorphism at P3 most likely eliminates the last two helices of CdeM in strain E10 (Fig. S3A and B).

### Supplemental References


Andersson, L. & M.W. Freeman, (1998) Functional changes in scavenger receptor binding conformation are induced by charge mutants spanning the entire collagen domain. *J Biol Chem* **273**: 19592-19601.

Bozue, J., K.L. Moody, C.K. Cote, B.G. Stiles, A.M. Friedlander, S.L. Welkos & M.L. Hale, (2007) Bacillus anthracis spores of the bclA mutant exhibit increased adherence to epithelial cells, fibroblasts, and endothelial cells but not to macrophages. *Infection and immunity* **75**: 4498-4505.

Brahmbhatt, T.N., B.K. Janes, E.S. Stibitz, S.C. Darnell, P. Sanz, S.B. Rasmussen & A.D. O'Brien, (2007) Bacillus anthracis exosporium protein BclA affects spore germination, interaction with extracellular matrix proteins, and hydrophobicity. *Infection and immunity* **75**: 5233-5239.

Calderon-Romero, P., P. Castro-Cordova, R. Reyes-Ramirez, M. Milano-Cespedes, E. Guerrero-Araya, M. Pizarro-Guajardo, V. Olguin-Araneda, F. Gil & D. Paredes-Sabja, (2018) Clostridium difficile exosporium cysteine-rich proteins are essential for the morphogenesis of the exosporium layer, spore resistance, and affect C. difficile pathogenesis. *PLoS Pathog* **14**: e1007199.

Costa, T., M. Serrano, L. Steil, U. Volker, C.P. Moran, Jr. & A.O. Henriques, (2007) The timing of cotE expression affects Bacillus subtilis spore coat morphology but not lysozyme resistance. *J Bacteriol* **189**: 2401-2410.

Cuff, J.A. & G.J. Barton, (1999) Evaluation and improvement of multiple sequence methods for protein secondary structure prediction. *Proteins* **34**: 508-519.

Darriba, D., G.L. Taboada, R. Doallo & D. Posada, (2011) ProtTest 3: fast selection of best-fit models of protein evolution. *Bioinformatics* **27**: 1164-1165.

Drozdetskiy, A., C. Cole, J. Procter & G.J. Barton, (2015) JPred4: a protein secondary structure prediction server. *Nucleic Acids Res* **43**: W389-394.

Escobar-Cortes, K., J. Barra-Carrasco & D. Paredes-Sabja, (2013) Proteases and sonication specifically remove the exosporium layer of spores of Clostridium difficile strain 630. *J Microbiol Methods* **93**: 25-31.

Finn, R.D., J. Clements, W. Arndt, B.L. Miller, T.J. Wheeler, F. Schreiber, A. Bateman & S.R. Eddy, (2015) HMMER web server: 2015 update. *Nucleic Acids Res* **43**: W30-38.

Finn, R.D., J. Clements & S.R. Eddy, (2011) HMMER web server: interactive sequence similarity searching. *Nucleic Acids Res* **39**: W29-37.

Heap, J.T., O.J. Pennington, S.T. Cartman & N.P. Minton, (2009) A modular system for Clostridium shuttle plasmids. *J Microbiol Methods* **78**: 79-85.

Hussain, H.A., A.P. Roberts & P. Mullany, (2005) Generation of an erythromycin-sensitive derivative of Clostridium difficile strain 630 (630Deltaerm) and demonstration that the conjugative transposon Tn916DeltaE enters the genome of this strain at multiple sites. *J Med Microbiol* **54**: 137-141.

Janoir, C., C. Deneve, S. Bouttier, F. Barbut, S. Hoys, L. Caleechum, D. Chapeton-Montes, F.C. Pereira, A.O. Henriques, A. Collignon, M. Monot & B. Dupuy, (2014) Adaptive strategies and pathogenesis of Clostridium difficile from in vivo transcriptomics. *Infection and immunity* **81**: 3757-3769.

Joshi, L.T., D.S. Phillips, C.F. Williams, A. Alyousef & L. Baillie, (2012) Contribution of spores to the ability of Clostridium difficile to adhere to surfaces. *Appl Environ Microbiol* **78**: 7671-7679.

Katoh, K. & D.M. Standley, (2013) MAFFT multiple sequence alignment software version 7: improvements in performance and usability. *Mol Biol Evol* **30**: 772-780.

Livingstone, C.D. & G.J. Barton, (1993) Protein sequence alignments: a strategy for the hierarchical analysis of residue conservation. *Comput Appl Biosci* **9**: 745-756.

Manganelli, R., R. Provvedi, C. Berneri, M.R. Oggioni & G. Pozzi, (1998) Insertion vectors for construction of recombinant conjugative transposons in Bacillus subtilis and Enterococcus faecalis. *FEMS Microbiol Lett* **168**: 259-268.

McKenney, P.T., A. Driks, H.A. Eskandarian, P. Grabowski, J. Guberman, K.H. Wang, Z. Gitai & P. Eichenberger, (2010) A distance-weighted interaction map reveals a previously uncharacterized layer of the Bacillus subtilis spore coat. *Curr Biol* **20**: 934-938.

Nickle, D.C., M. Rolland, M.A. Jensen, S.L. Pond, W. Deng, M. Seligman, D. Heckerman, J.I. Mullins & N. Jojic, (2007) Coping with viral diversity in HIV vaccine design. *PLoS Comput Biol* **3**: e75.

Paredes-Sabja, D. & M.R. Sarker, (2012) Adherence of Clostridium difficile spores to Caco-2 cells in culture. *J Med Microbiol* **61**: 1208-1218.

Pereira, F.C., L. Saujet, A.R. Tome, M. Serrano, M. Monot, E. Couture-Tosi, I. Martin-Verstraete, B. Dupuy & A.O. Henriques, (2013) The spore differentiation pathway in the enteric pathogen Clostridium difficile. *PLoS Genet* **9**: e1003782.

Pizarro-Guajardo, M., P. Calderon-Romero, P. Castro-Cordova, P. Mora-Uribe & D. Paredes-Sabja, (2016) Ultrastructural Variability of the Exosporium Layer of Clostridium difficile Spores. *Appl Environ Microbiol* **82**: 2202-2209.

Rabi, R., L. Turnbull, C.B. Whitchurch, M. Awad & D. Lyras, (2017) Structural Characterization of Clostridium sordellii Spores of Diverse Human, Animal, and Environmental Origin and Comparison to Clostridium difficile Spores. *mSphere* **2**.

Rosenberg, M., (1984) Bacterial adherence to hydrocarbons: a useful technique for studying cell surface hydrophobicity. *FEMS Microbiology Letters* **22**: 289-295.

Saujet, L., M. Monot, B. Dupuy, O. Soutourina & I. Martin-Verstraete, (2011) The key sigma factor of transition phase, SigH, controls sporulation, metabolism, and virulence factor expression in Clostridium difficile. *J Bacteriol* **193**: 3186-3196.

Saujet, L., F.C. Pereira, M. Serrano, O. Soutourina, M. Monot, P.V. Shelyakin, M.S. Gelfand, B. Dupuy, A.O. Henriques & I. Martin-Verstraete, (2013) Genome-wide analysis of cell type-specific gene transcription during spore formation in Clostridium difficile. *PLoS Genet* **9**: e1003756.

Stabler, R.A., M. He, L. Dawson, M. Martin, E. Valiente, C. Corton, T.D. Lawley, M. Sebaihia, M.A. Quail, G. Rose, D.N. Gerding, M. Gibert, M.R. Popoff, J. Parkhill, G. Dougan & B.W. Wren, (2009) Comparative genome and phenotypic analysis of Clostridium difficile 027 strains provides insight into the evolution of a hypervirulent bacterium. *Genome Biol* **10**: R102.

Stamatakis, A., (2014) RAxML version 8: a tool for phylogenetic analysis and post-analysis of large phylogenies. *Bioinformatics* **30**: 1312-1313.

Waterhouse, A.M., J.B. Procter, D.M. Martin, M. Clamp & G.J. Barton, (2009) Jalview Version 2--a multiple sequence alignment editor and analysis workbench. *Bioinformatics* **25**: 1189-1191.

Zheng, L.B., W.P. Donovan, P.C. Fitz-James & R. Losick, (1988) Gene encoding a morphogenic protein required in the assembly of the outer coat of the Bacillus subtilis endospore. *Genes Dev* **2**: 1047-1054.

### Supplemental Figure Legends

**Figure S1 – Use of the SNAP substrate TMR-Star to monitor the expression of *cdeM* during spore morphogenesis**. **A**: samples were collected from 24 h liquid SM cultures of strains expressing transcriptional $P_{cotE}$-$SNAP^{Cd}$, $P_{cdeC}$-$SNAP^{Cd}$ , and $P_{cdeM}$-$SNAP^{Cd}$ fusions and the cells labeled with the SNAP substrate TMR-Star substrate and the membrane dye MTG. The cells were examined by phase contrast and fluorescence microscopy and scored for expression of the different fusions. The numbers are the percentage of sporangia showing expression of the various fusions at the represented stages in morphogenesis, defined as in the legend for figure 1A, except that *a*, asymmetric septation, is not represented; *b*, an intermediate stage in engulfment; *c*, engulfment completion; *d*, phase grey spores; *e*, phase bright spores (with the various spore layers represented); *f*, free spores. The vertical grey arrows represent the window of expression of the indicated transcriptional $SNAP^{Cd}$ fusions. **B**: the $P_{cdeM}$-$SNAP^{Cd}$ fusion was introduced in a *sigK* insertional mutant and expression monitored as described above. The image shows sporangia at late stages in morphogenesis, when the phenotype characteristic of a *sigK* mutant, *i.e.*, the formation of partially refractile spores, is seen. Scale bar in A and B, 1 µm.

**Figure S2** - **Complementation of *cdeM::ermB* using transposon Tn*916***. **A**: Schematic representation of the integration of plasmid pFT71 (pSMB47 pseudo-suicide vector containing *cdeM* promoter region and coding sequence) into Tn*916*. The main features of pFT71 plasmid (on top) are indicated, together with the position of the SalI/HindIII sites used to clone *cdeM*. The plasmid integrates into Tn916, initially present in *B. subtilis* BS49 chromosome, via a single crossover homologous recombination event, represented by a cross, and that occurs downstream of *orf14*.

Tn*916* containing pFT71 was then transferred from BS49 to *C. difficile cdeM::ermB* chromosome via bacterial conjugation (bottom). **B**: Chromosomal DNA of *B. subtilis* BS49 transformants or of *C. difficile cdeM::ermB* conjugants was screened by PCR using primer pairs cdeMFw/ermR to confirm the presence of pFT71, with PcdeMFw/cdeMR to confirm the presence of second intact copy of *cdeM* in the genome of the *cdeM::ermB* mutants, and with PtcdBFw/Rev primers to confirm that the transconjugants were *C.difficile*, and not *B.subtilis* used for conjugation and that was able to grow in the anaerobic environment (left panel). Integration of pFT71 in Tn*916* at the expected place, as depicted in A, was also confirmed by screening chromosomal DNA of *B. subtilis* BS49 transformants and of *C. difficile cdeM::ermB* conjugants with primer pair PcdeMR/orf13R (right panel). The position of the DNA size markers is indicated on the left side of the panel.

**Figure S3** - **CdeM is a polymorphic cysteine rich protein that undergoes multimerization *in vitro***. **A**: Aminoacid sequence alignment of the cysteine-rich CdeM coded for by the genomes of thirteen different *C. difficile* strains (*Ensemble Bacteria* database). The genomes represent the thirteen unique amino acid sequences among the 229 homologous sequences that we recovered from the database (see the Supplemental Material for details). The sequences were aligned using MAFFT and visualized using JalView (Waterhouse *et al.*, 2009). The amino acid residues (in single letter notation) are colored according to the default Clustal X color scheme (www.ebi.org). The panels below the alignment are statistics that were estimated using available tools on Jalview ((Waterhouse *et al.*, 2009); see also the Supplemental Material for a description). Note that four regions of polymorphism are seen, labeled P1 through P4; corresponding to insertions/deletions (P1, P3 and P4; shown by the blue dots and arrows in the corresponding position of the CdeM protein from strain

630) and amino acid substitutions (P2, red dot and arrow). **B**: Schematic representation of the organization predicted for the CdeM protein, according to the alignment in **A**: residues 1-37 form an N-terminal variable region; residues 37-130 form a long $\alpha$-helical domain; residues 130-160 form a C-terminal Cys-rich domain. **C**: a CdeM-His$_6$ fusion was overproduced in *E. coli*, partially purified by $Ni^{2+}$-affinity chromatography and the protein resolved by SDS-PAGE (on a15% gel) in the presence or in the absence of DTT (10 mM). Coomassie stained bands present in the brace area correspond to multimeric forms of CdeM-His$_6$. The black arrow points to an abundant multimeric of CdeM-His$_6$, possibly a trimer, that migrates at around 60 kDa, and the red arrow indicates the position of monomeric CdeM-His$_6$, in the 20 kDa region of the gel (the predicted mass of the protein is 19.22 kDa). The position of molecular weight (MW) markers (in kDa) is shown on the left side of the panel.

**Figure S4 – CdeM is exposed at the spore surface**. **A**: Spore-associated CdeM is accessible to trypsin. Proteins were extracted from WT spores untreated (-) or following treatment (+) with trypsin. Spores from *cdeM*::*ermB* were also treated with trypsin and the proteins resolved by SDS-PAGE. In both A and B, the position of monomeric (black arrows) and multimeric forms of CdeM (red arrows) is indicated; the blue arrow shows a possible degradation product of CdeM. The position of molecular weight (MW) markers (in kDa) is shown on the left side of panels A and B. **B**: Spores produced from the WT or the *cdeM*::*ermB* congenic mutant were purified and subject to immunofluorescence microscopy with an anti-CdeM antibody and a secondary antibody conjugated to Alexa 488, without or following treatment with trypsin. The white arrows show the localization of the fluorescence signal in selected spores. Scale bar, 1 µm. **C**: CdeM is not released by washing spores with a concentrated solution of KCl. Coat proteins were extracted from spores not washed (-) or washed (+) with 1 M

KCl, and resolved by SDS-PAGE. The supernatant fraction (S) of the spore wash was also analyzed by SDS-PAGE. The gel was stained with Coomassie. **D**: *cdeM* mutant spores are less hydrophobic than WT spores. Hydrophobicity of density gradient-purified spores of the wild type, the *cdeM::ermB* mutant and the complementation strain was estimated by a water/hexadecane partitioning assay as described in the Materials and Methods. Error bars are from 3 independent experiments.

**Figure S5 – Appendage characterization in spores of strain R20291. A:** quantification of the percentage of type A and type B spores (as defined in Fig. 4A; see also the main text), in a population of R20291 spores. **B**: Box plots showing the median, lower and upper quartiles of the distance measured (in µm), for each of the indicated strains, in the phase contrast or FM4-64 images. Each point is a measurement of a single spore. Differences between the two strains are statistically significant for all measures ($P < 0.001$; Welch t-test for width and length FM4-64; Wilcoxon rank sum test for length Ph. contrast). **C:** representative shells isolated from spores of the 630Δ*erm*, R20291 and *cdeM*::*ermB* mutant (background of 630Δ*erm*). Sclae bar, 1 µm. **D:** distribution of the shell thickness measurements obtained for spores of 630Δ*erm* (green symbols), R20291 (red), and the *cdeM*::*ermB* mutant.

**Figure S6 – Phase contrast microscopy of WT (630Δ*erm*) spores fractionated on gradients of metrizoic acid**. Phase contrast microscopy of WT and *cdeM::ermB* spores fractionated on step gradients of Gastrografin. Samples are from the 45%, 50% and 55% layers. Polar appendages are present in WT and *cdeM::ermB* spores (red arrows) but are often shorter and disorganized in the *cdeM* mutant (yellow arrows). The blue arrow points to material that may be composed of aggregated empty shells. Scale bar, 1 µm.

**Figure S7 – Distribution of CdeM types**. **A**: *cdeM* gene tree inferred with a maximum likelihood on the protein sequences using the HIVb+G model of protein evolution (Nickle *et al.*, 2007). This group of sequences represent all of the significant hits (E-value threshold, $10^{-4}$) that we obtained by querying the sequence of CdeM from strain 630 against the Ensemble Bacteria database using *phmmer* (Finn *et al.*, 2015, Finn *et al.*, 2011). The tree was inferred using RaxML (Stamatakis, 2014). Numbers on the nodes are bootstrap support (only bootstrap values higher than 50 are shown). The tree is drawn to scale, and the scale bar indicates expected number of amino acid substitutions per site. The numbers at the end of the strain names indicate the number of genomes, from a total of 229 genomes, that were identified with highly similar protein sequences. The sequences shown are the subset that were selected to represent the polymorphism present in this protein (see also figure S3). The three more represented types of sequences are highlighted by the dots (strains 630, E24 and R20291, types A, E and D, respectively; see also Table S3).

### Supplemental Tables

#### Table S1 - Bacterial strains and plasmids.

| Strain/ Plasmid | Relevant Genotype/Properties* | Origin/ Reference |
|---|---|---|
| **STRAINS** | | |
| ***E. coli*** | | |
| DH5α | F– Φ80*lac*ZΔM15 Δ(*lac*ZYA-*arg*F) U169 *rec*A1 *end*A1 *hsd*R17 (rK–, mK+) *pho*A *sup*E44 λ– *thi*-1 *gyr*A96 *rel*A1 | Invitrogen |
| BL21 (DE3) | *fhuA2 [lon] ompT gal (λ DE3) [dcm] ΔhsdS* *λ DE3 = λ sBamHIo ΔEcoRI-B int::(lacI::PlacUV5::T7 gene1) i21 Δnin5* | Novagen |
| HB101 (RP4) | F$^-$ *λ*$^-$ *araC14 leuB6*(Am) DE(*gpt-proA*)62 *lacY1 glnX44*(AS) *galK2* (Oc) *recA13 rpsL20*(StrR) *xylA5 mtl*-1 *thiE1* *hsdS*20($r_B^-$, $m_B^-$); carries conjugational plasmid pRP4. | Laboratory stock |
| AHCD070 | BL21 (DE3)/ pFT41 | This work |
| AHCD076 | HB101 (RP4)/ pMTL84121 | " |
| AHCD113 | HB101 (RP4)/ pFT52 | " |
| AHCD127 | HB101 (RP4)/ pFT57 | " |
| ***B. subtilis*** | | |
| BS49 | CU2189::Tn*916* | P. Mullany |
| AHCD159 | BS49 Tn*916*::pFT71 | This work |
| ***C. difficile*** | | |
| 630Δ*erm* | *C. difficile* 630Δ*erm*, ribotype 012 | (Hussain *et al.*, 2005) |
| R20291 | Epidemic strain, ribotype 027 | (Stabler *et al.*, 2009) |
| AHCD532 | 630Δ*erm sigE*::intron *ermB* | (Pereira *et al.*, 2013) |
| AHCD533 | 630Δ*erm sigF*::intron *ermB* | " |
| AHCD534 | 630Δ*erm sigG*::intron *ermB* | " |
| AHCD535 | 630Δ*erm sigK*::intron *ermB* | " |
| 630Δ*erm M* | 630Δ*erm cdeM*::intron *ermB* | (Janoir *et al.*, 2014) |
| AHCD543 | 630Δ*erm* containing pMTL84121 | This work |
| AHCD599 | 630Δ*erm* containing pFT52 | " |
| AHCD620 | AHCD535 containing pFT52 | " |
| AHCD715 | 630Δ*erm cdeM* containing pFT71 | " |
| AHCD871 | 630Δ*erm* containing pCAF8 ($P_{cdeC}$-*SNAP*$^{Cd}$) | " |
| AHCD824 | 630Δ*erm* containing pCAF5 ($P_{cdeM}$-*SNAP*$^{Cd}$) | " |
| AHCD620 | 630Δ*erm* containing pCAF7 ($P_{cotE}$-*SNAP*$^{Cd}$) | " |
| **PLASMIDS** | | |
| pMTL84121 | *Clostridium* modular plasmid containing *catP* (Cm$^R$/Tm$^R$) | (Heap *et al.*, 2009) |
| pET33b | His-tag fusion protein expression vector (Km$^R$) | Novagen |
| pFT47 | pMTL84121-*SNAP*$^{Cd}$ (Cm$^R$/Tm$^R$) | (Pereira *et al.*, 2013) |
| pSMB47 | Tn*916* integrational vector (Cm$^R$/Tm$^R$) | (Manganelli *et al.*, 1998 |
| pFT41 | *cdeM* in pET33b (Km$^R$) | This work |
| pFT43 | pMTL84121 containing 483-bp of the *cdeM* coding region and 534-bp upstream of the coding region region | " |
| pFT52 | pFT47 containing the *cdeM* promoter region (P*cdeM-SNAP*$^{Cd}$) | " |
| pFT57 | CdeM-SNAP in pMTL84121 | " |
| pFT71 | *cdeM* coding region and 534-bp upstream promoter region from pFT43 cloned as SalI/HindIII fragment in pSMB47 | " |

| | | |
|---|---|---|
| pCAF5 | $P_{cdeM}$-*SNAP*$^{Cd}$ in pMTL84121 | “ |
| pCAF7 | $P_{cotE}$-*SNAP*$^{Cd}$ in pMTL84121 | “ |
| pCAF8 | $P_{cdeC}$-*SNAP*$^{Cd}$ in pMTL84121 | “ |

$^{*}$Cm$^{R}$, cloranfenicol resistance; TM$^{R}$, thianfenicol resistance; Km$^{R}$, kanamycin resistance; *ermG*, erythromycin resistance.

#### Table S2 – Oligonucleotides used in this study.

| Primer | Sequence (5´to 3´)[a] | Description |
|---|---|---|
| DNApolIII-F | TCCATCTATTGCAGGGTGGT | qRT-PCR, *dnaF* (CD1305 |
| DNApolIII-R | CCCAACTCTTCGCTAAGCAC | " |
| IMV552 | CCAGGAGCAAAAGAGGCTTTA | qRT-PCR, *sspA* (CD2688 |
| IMV553 | TCCAGCCATTTGTTGTTCAG | " |
| LS100 | CACCACCTCAATGTGGAAAA | qRT-PCR, *spoIIIAA* (CD1192) |
| LS101 | GCTCCTGCTATCTCATTACGC | " |
| LS156b | GCAGTAGGTGGAGTTTGGTTT | qRT-PCR, *spoIIQ* (CD012 |
| LS157 | TGAATCCGTTGCAGTAGGG | " |
| cdeM-pET33b Fw | GATATGTTCCATGGAAAATAAAAAATGTTATTC | 5'-*cdeM* NcoI |
| cdeM-pET33b Rev | CTAATTCTCGAGTTTTCTACAGCAGTTACA | 3'-*cdeM* XhoI-no STOP |
| PcdeM Fw | GCTGTCGACGCTAGAATAGACAATGAG | 5'-promoter *cdeM* SalI |
| PcdeM Rev | TGCAGCCTCGAGTTATATAACATATCTCTCCC | 3-promoter *cdeM* XhoI |
| cdeM Rev | CAATTACTCGAGTTATTTTCTACAGCAGTT | 3'-*cdeM* XhoI |
| SNAP-cdeM Linker Fw | GGTGGTGGTGGATCCGCAGCTGCTGATAAAGATTGT AATGAAG | 5'- $SNAP^{Cd}$ -linker |
| cdeM Linker Rev | GGATCCACCACCACCAAGTTTTCTACAGCAGTTACA | 3'-*cdeM*-linker |
| SNAPtag BamHI_Rev | GCTGTTGGATCCAAGCTTTCCTTACCC | 3'-$SNAP^{Cd-}$BamHI |
| ErmRAM-R | ACGCGTGCGACTCATAGAATTATTTCCTCCCG | PCR *erm* pSMB47 |
| Orf13 Rev | GAGAGCTTGAAGTTCATCTGTCAG | PCR *orf13* Tn*916* |

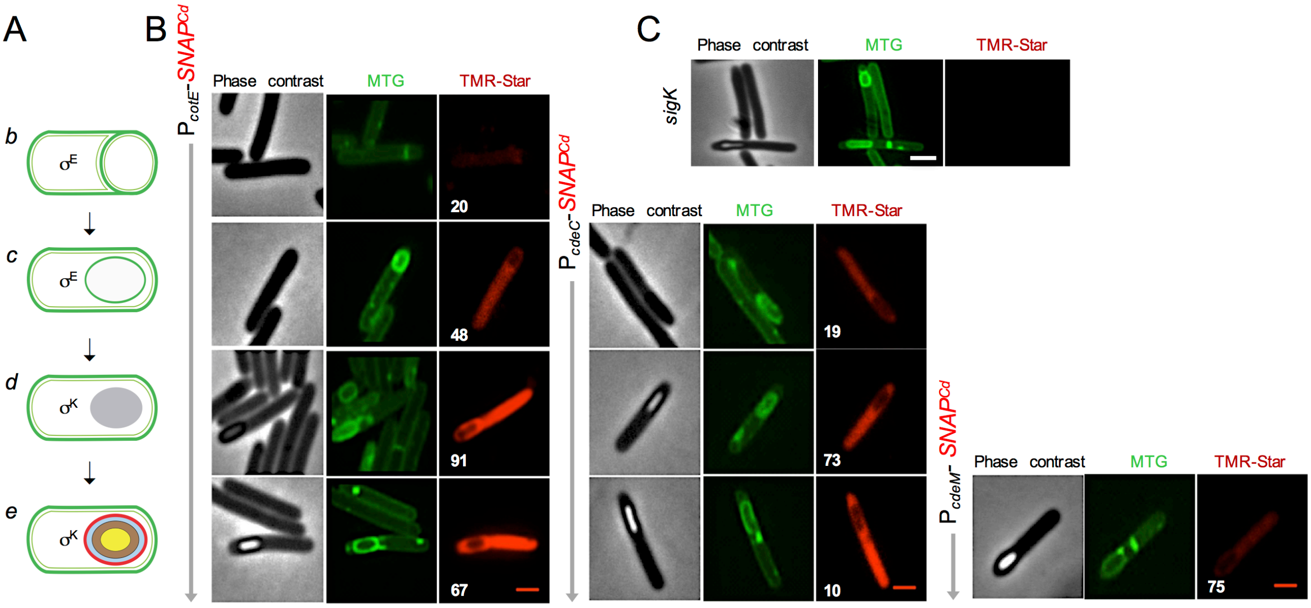
A
B
C
b
σE
c
σE
d
σK
e
σK
PcotE-SNAPCd
Phase contrast
MTG
TMR-Star
20
48
91
67
sigK
PcdeC-SNAPCd
19
73
10
PcdeM-SNAPCd
75

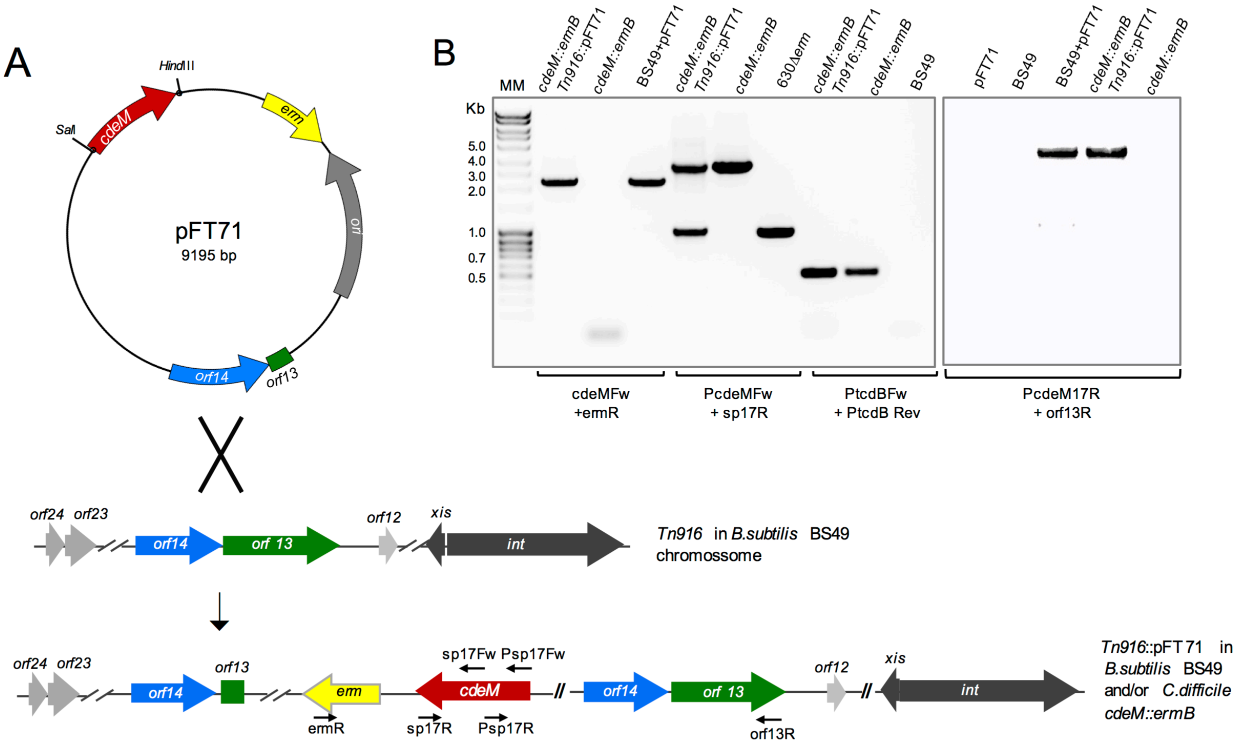
A
B
HindIII
SalI
cdeM
erm
ori
pFT71
9195 bp
orf14
orf13
MM
Kb
5.0
4.0
3.0
2.0
1.0
0.7
0.5
cdeM::ermB
Tn916::pFT71
cdeM::ermB
BS49+pFT71
cdeM::ermB
Tn916::pFT71
cdeM::ermB
630Δerm
cdeM::ermB
Tn916::pFT71
cdeM::ermB
BS49
pFT71
BS49
BS49+pFT71
cdeM::ermB
Tn916::pFT71
cdeM::ermB
cdeMFw
+ermR
PcdeMFw
+ sp17R
PtcdBFw
+ PtcdB Rev
PcdeM17R
+ orf13R
orf24 orf23
orf14
orf 13
orf12
xis
int
Tn916 in B.subtilis BS49
chromossome
orf24 orf23
orf14
orf13
erm
cdeM
sp17Fw Psp17Fw
ermR
sp17R
Psp17R
orf14
orf 13
orf13R
orf12
xis
int
Tn916::pFT71 in
B.subtilis BS49
and/or C.difficile
cdeM::ermB

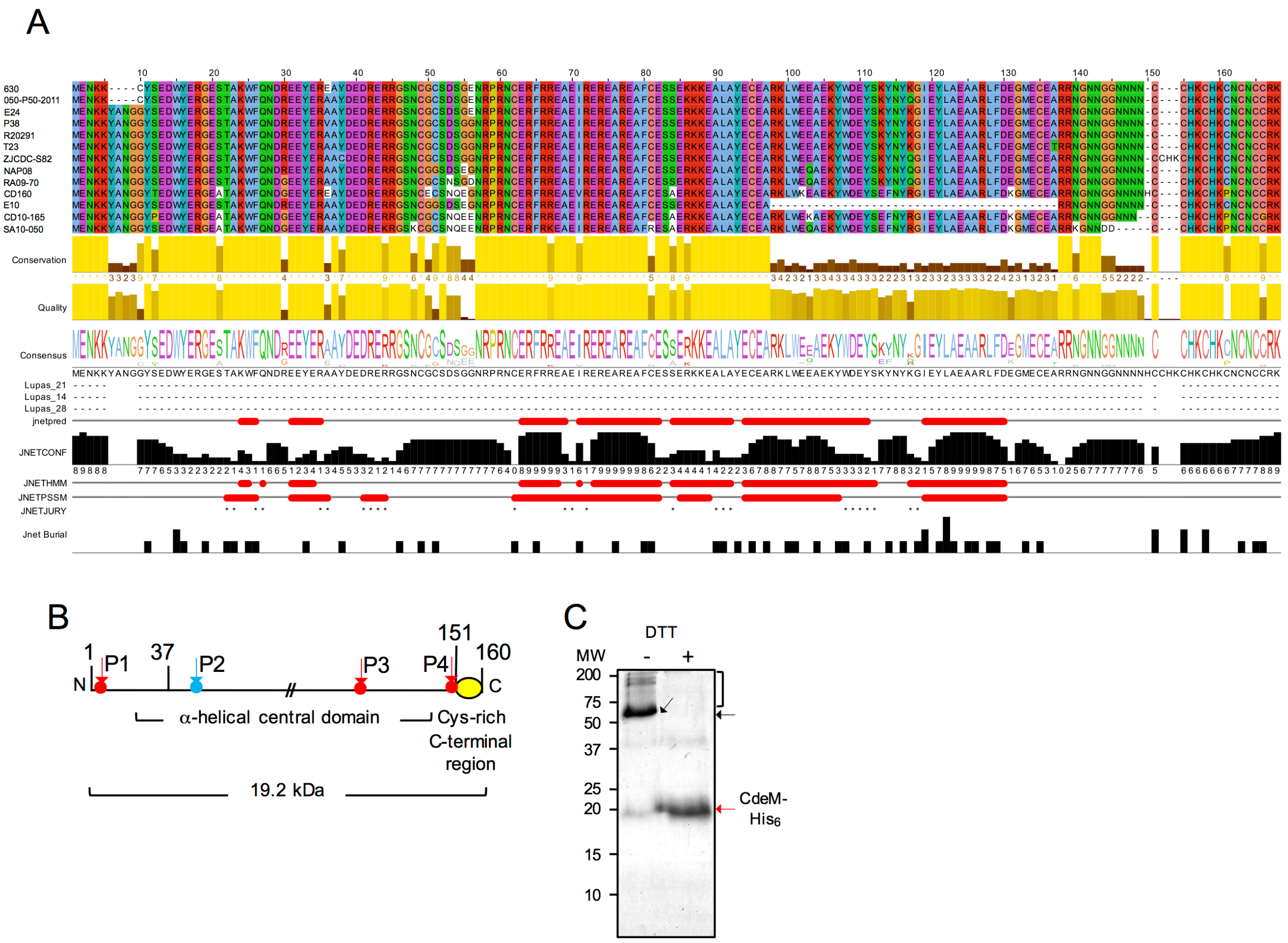

A
630
050-P50-2011
E24
P38
R20291
T23
ZJCDC-S82
NAP08
RA09-70
CD160
E10
CD10-165
SA10-050
Conservation
Quality
Consensus
Lupas_21
Lupas_14
Lupas_28
jnetpred
JNETCONF
JNETHMM
JNETPSSM
JNETJURY
Jnet Burial
B
1
37
151
160
N
C
P1
P2
P3
P4
α-helical central domain
Cys-rich C-terminal region
19.2 kDa
C
DTT
MW
-
+
200
75
50
37
25
20
15
10
CdeM-His6

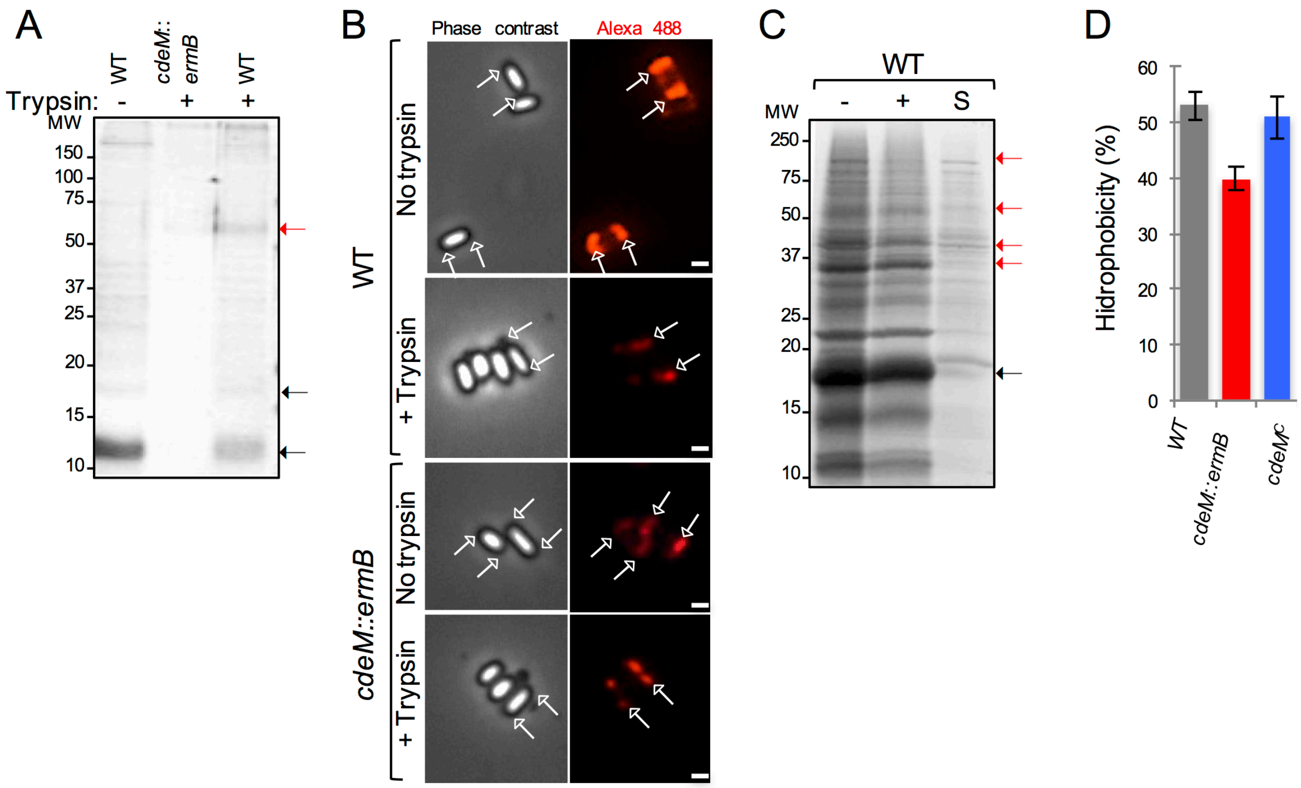

A
WT
cdeM::
ermB
WT
Trypsin: - + +
MW
150
100
75
50
37
25
20
15
10
B
Phase contrast
Alexa 488
WT
No trypsin
+ Trypsin
cdeM::ermB
No trypsin
+ Trypsin
C
WT
MW
- + S
250
75
50
37
25
20
15
10
D
Hidrophobicity (%)
60
50
40
30
20
10
0
WT
cdeM::ermB
cdeMC

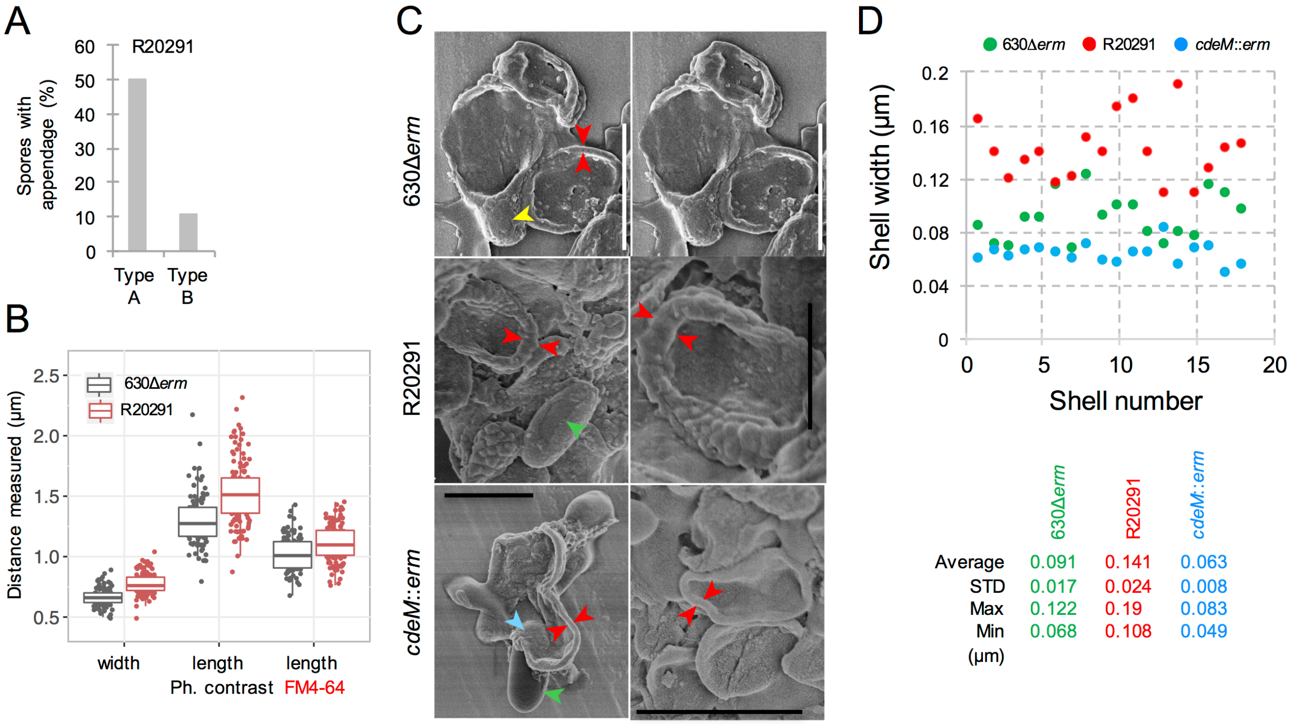
A
R20291
Spores with appendage (%)
60
50
40
30
20
10
0
Type A
Type B
B
Distance measured (µm)
2.5
2.0
1.5
1.0
0.5
630Δerm
R20291
width
length
length
Ph. contrast
FM4-64
C
630Δerm
R20291
cdeM::erm
D
630Δerm
R20291
cdeM::erm
Shell width (µm)
0.2
0.16
0.12
0.08
0.04
0
0
5
10
15
20
Shell number
630Δerm R20291 cdeM::erm
Average 0.091 0.141 0.063
STD 0.017 0.024 0.008
Max 0.122 0.19 0.083
Min 0.068 0.108 0.049
(µm)

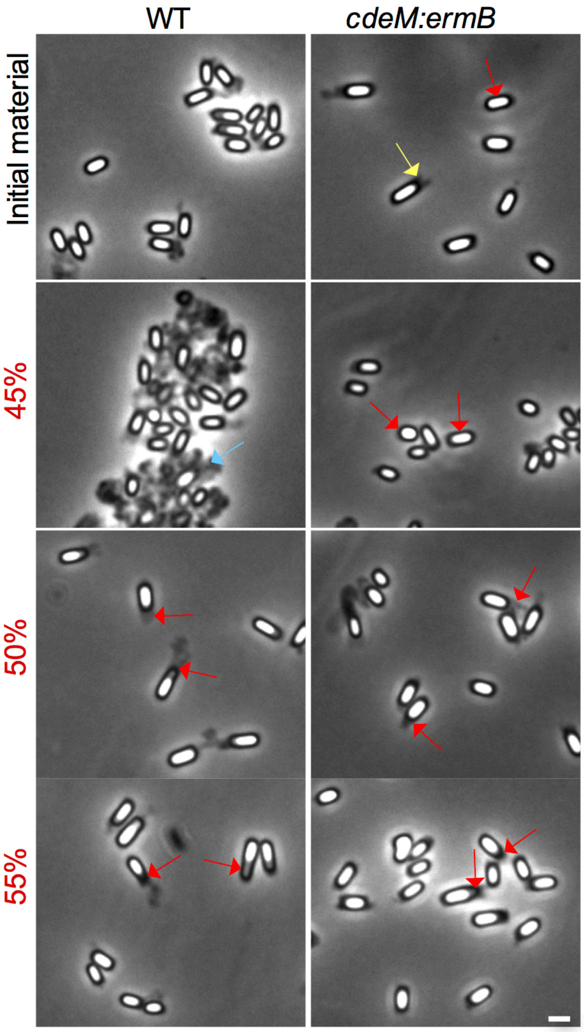

WT
cdeM:ermB
Initial material
45%
50%
55%

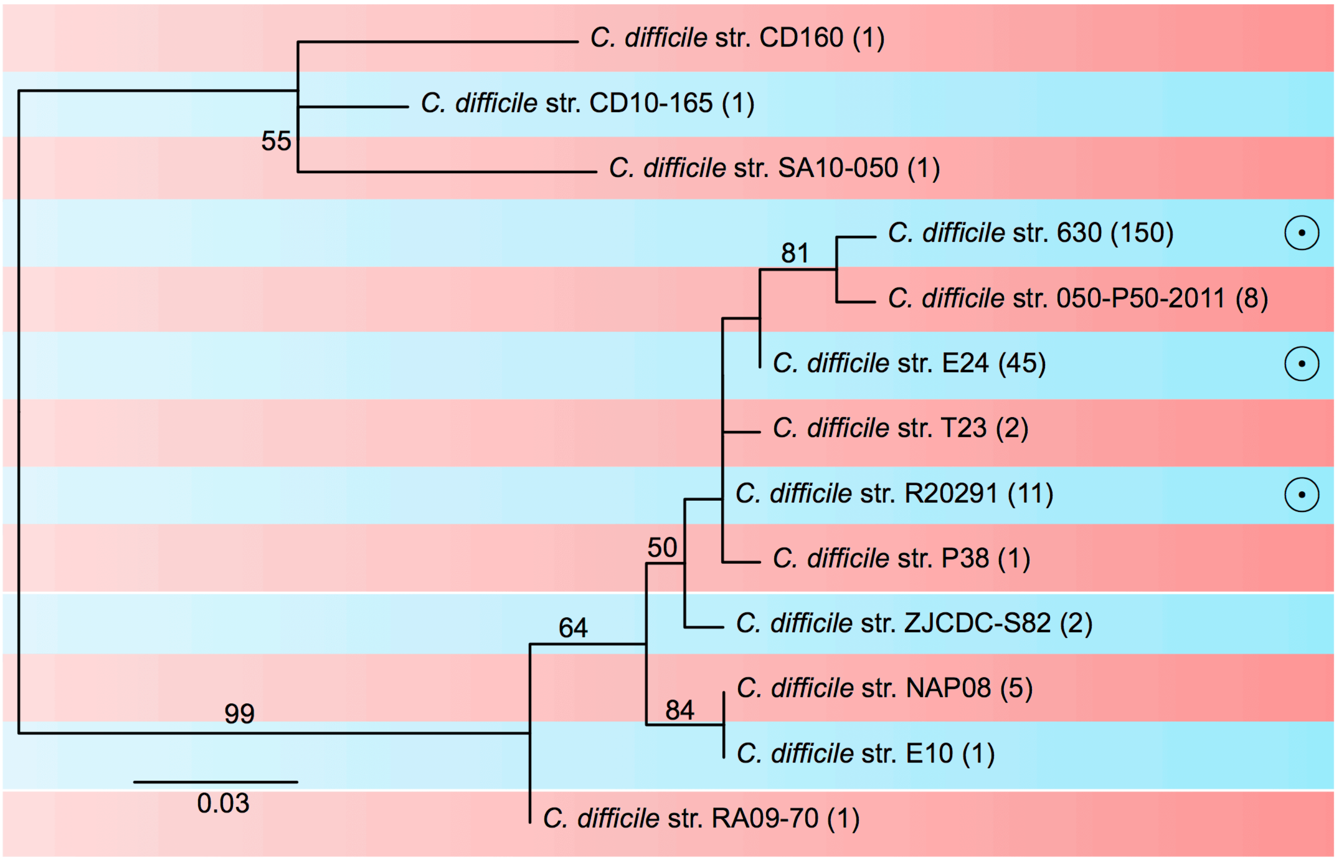

C. difficile str. CD160 (1)
C. difficile str. CD10-165 (1)
55
C. difficile str. SA10-050 (1)
C. difficile str. 630 (150)
81
C. difficile str. 050-P50-2011 (8)
C. difficile str. E24 (45)
C. difficile str. T23 (2)
C. difficile str. R20291 (11)
50
C. difficile str. P38 (1)
64
C. difficile str. ZJCDC-S82 (2)
C. difficile str. NAP08 (5)
99
84
C. difficile str. E10 (1)
0.03
C. difficile str. RA09-70 (1)
